# A deep learning framework for building INDEL mutation rate maps

**DOI:** 10.1101/2025.11.18.689146

**Authors:** Shuyi Deng, Hui Song, Cai Li

## Abstract

Germline short insertions and deletions (INDELs) are pervasive genetic variants that shape genome evolution and contribute to human disease. However, accurately quantifying fine-scale INDEL mutation rates remains challenging due to data limitations and the diversity of INDEL subtypes. Here, we present MuRaL-indel, a deep learning framework that predicts germline INDEL mutation rates by leveraging long-range sequence context through a U-Net architecture. Using extensive rare variant data from large population cohorts, MuRaL-indel generates base-resolution, length-specific mutation rate maps for the human genome and achieves superior accuracy compared with existing models across multiple genomic scales. We successfully apply MuRaL-indel to three non-human species (*Macaca mulatta*, *Drosophila melanogaster*, and *Arabidopsis thaliana*), demonstrating its broad applicability across taxa. Using the predicted mutation rate maps, we reveal the mutational landscape around human coding genes and show that MuRaL-indel–derived constraint scores better prioritize pathogenic INDELs than previous models. Through deep learning interpretability analyses, we uncovered sequence motifs—including both repeat and non-repeat elements—associated with elevated INDEL mutability, providing insights into underlying mutational mechanisms. Together, MuRaL-indel establishes a generalizable and scalable framework for building high-resolution INDEL mutation rate maps, offering a valuable resource for studies of genome evolution, mutational mechanism, variant interpretation, and genetic disease.

## Introduction

Germline mutations occur in germ cells and during zygote formation, serving as an ultimate source of genetic diversity^1^. Alongside single nucleotide variants (SNVs), small insertion/deletion (INDEL) mutations represent another common type of genetic variations, accounting for over 10% of small variants in human populations^2^. Although INDELs occur less frequently in the germline than SNVs, many of them are located in functionally important sites and thus potentially influence organismal traits and disease. Notably, non-triplet INDELs in coding regions (known as frameshift mutations) can disrupt open reading frames, leading to likely unfunctional or damaging proteins^3^. Germline INDEL mutations have been implicated in various human diseases such as cystic fibrosis^4^ and hemophilia^5^. The estimated germline INDEL mutation rate in the human genome ranges from 1.16 × 10^-9^ to 2.5 × 10^-9^ per generation per base pair^6^. Similar to point mutations, INDEL mutation rates show pronounced heterogeneity across multiple genomic scales, ranging from short sequence motifs^7,8^ to chromosomal regions^9,10^. Accurate quantification of INDEL mutation rates is critical for many analyses, such as reconstructing ancestral sequence^11,12^, estimating selection constraints^8,13^ and identifying disease-associated mutations^14^.

Building a fine-scale INDEL mutation rate map currently faces substantial obstacles. Firstly, compiling a high-quality INDEL dataset for mutation rate modeling is difficult. Early studies predominantly inferred INDELs through cross-species genomic sequence alignment^10,15–18^, yielding data that has been subject to prolonged natural selection and thus fails to directly reflect the intrinsic patterns of germline INDEL mutagenesis. Even with the advent of large-scale trio-based population sequencing projects, the number of *de novo* INDEL mutations identified remains insufficient to support effective fine-scale modeling. Recent studies suggested that extremely rare variants observed in population genomic data may serve as proxy approximations for *de novo* mutations^19^, offering a practical approach to acquiring a large number of training variants. Secondly, many factors have been reported to be associated with INDEL mutagenesis^20^ and how to model mutation rates with related factors is challenging. Previous research has attempted to predict INDEL mutation rates using proximal nucleotide sequence features^14,18,21^ or large-scale genomic functional annotations^10^ (e.g., DNA methylation, replication timing and recombination rate). However, the resulting mutation rate estimates remained suboptimal.

Existing INDEL mutation rate prediction methods suffer from several critical limitations. Firstly, many approaches rely on INDEL mutation data derived from cross-species sequence alignments or population-wide polymorphic sites^10,14–18^, which have been heavily affected by selection. Secondly, most methods are limited to incorporating only very short sequence context^18,21^ or modeling a single mutational mechanism^14^, thereby failing to comprehensively characterize the multifaceted mutational landscape of INDELs. The relationship between longer sequence features and mutations remains underexplored in these models. Thirdly, many existing frameworks employ simplistic linear or generalized linear models^10,14,18^ to describe the relationship between mutational features and mutation rates. Such models may fail to capture the complex combinatorial effects of genomic context and mutational processes. Finally, traditional predictive models for estimating INDEL mutation rates typically neglect the INDEL length, which is a key factor governing the magnitude of mutational impact. Developing methods to quantify the site-specific probabilities of INDELs of varying lengths would enhance our understanding of mutation rate heterogeneity and enable more sophisticated downstream analyses, such as INDEL pathogenicity assessment in coding genes.

Deep learning approaches have demonstrated remarkable performance in addressing complex predictive problems and are increasingly being applied to genomic studies^22–24^. We previously developed a deep learning model (named MuRaL) for predicting base-resolution SNV mutation rates, which showed superior performance compared to traditional methods^25^. However, the distinct mutation types and underlying mechanisms of SNVs and INDELs preclude the direct application of the MuRaL model to INDEL rate prediction. Like SNVs, INDEL mutation rates are strongly influenced by local sequence context^7,18^, and many functional genomic features are sequence-dependent. We therefore propose that deep learning could offer a promising avenue to capture the complex mutability signals embedded in genomic sequences for accurate INDEL mutation rate prediction.

Motivated by these considerations, we developed MuRaL-indel (Mutation Rate Learner for INDELs), an extension of the MuRaL framework^25^ tailored for germline INDEL mutation rate prediction. This model estimates base-resolution INDEL mutation rates for specific length classes across the genome using DNA sequence as its sole input. Comprehensive evaluation using human INDEL variant datasets demonstrates that MuRaL-indel predictions are highly correlated with empirically observed mutation rates across various scales ranging from kilobases to megabases. Compared to current state-of-the-art models, MuRaL-indel achieves superior performance, with substantial improvements in predictive accuracy. Furthermore, we show that MuRaL-indel is readily adaptable to non-human species. We also demonstrate that the improved mutation rate estimates empower practical applications such as prioritizing pathogenic mutations and uncovering sequence determinants of INDEL mutability.

## Results

### Design of the MuRaL-indel model

Similar to the kmerPaPa model by Bethune et al.^21^, we focused on predicting the probabilities of insertions and deletions at potential genomic breakpoint sites. Specifically, each insertion is associated with a single breakpoint, while each deletion has two breakpoints—designated as the “deletion start” and “deletion end” (**Fig. 1a**). For each such breakpoint, our goal is to predict mutation probabilities of INDELs of specific lengths (ignoring the inserted/deleted DNA content) based on the upstream and downstream sequence context of the breakpoint. A key distinction from kmerPaPa is that their model provides only an overall mutation rate at each breakpoint, without resolving mutation probabilities by INDEL length.

**Fig. 1.**
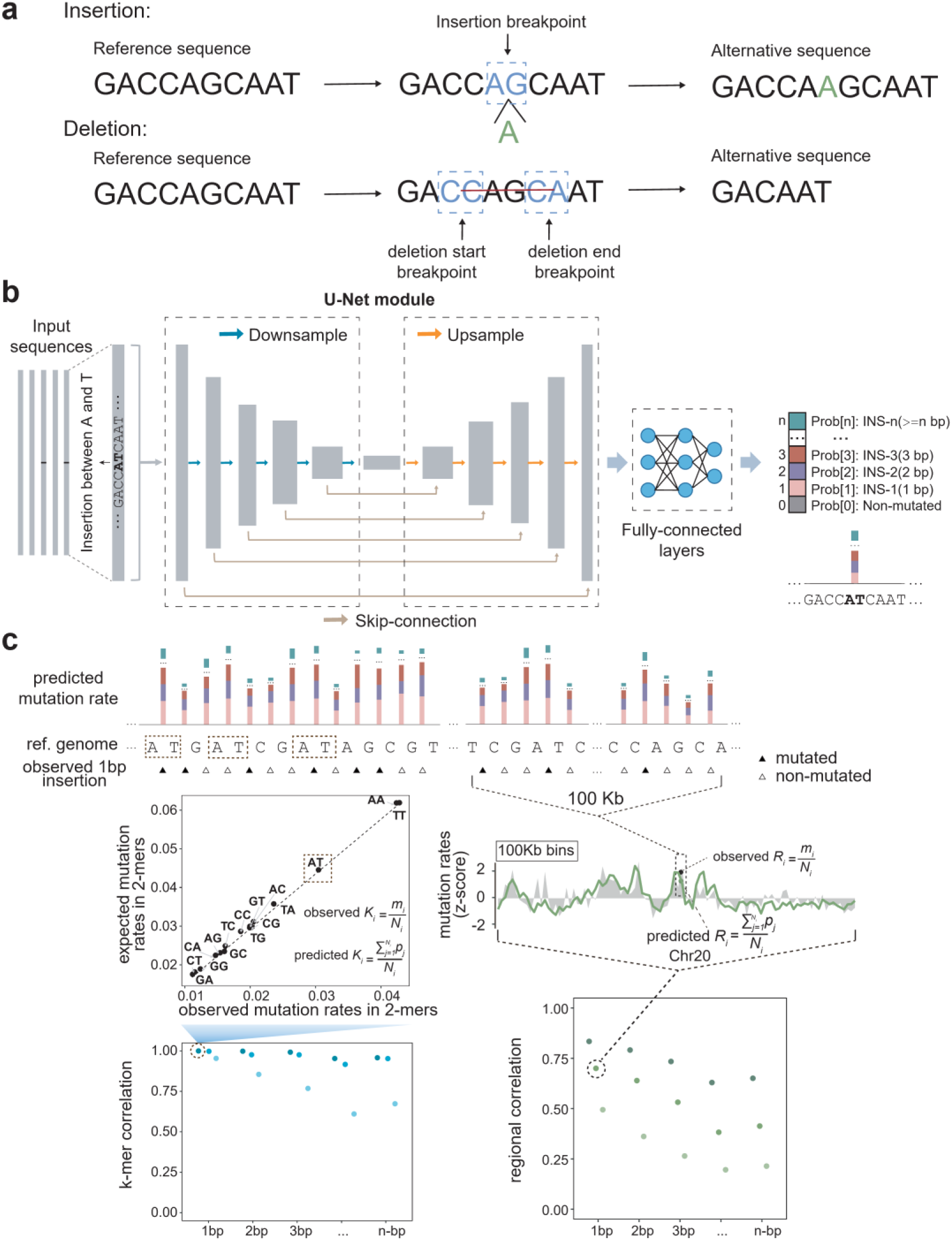
Overview of the MuRaL-indel model and evaluation metrics. (**a**) Illustration of INDEL breakpoints. Each insertion mutation occurs at a single breakpoint, while each deletion mutation spans two breakpoints—designated as “deletion start” and “deletion end”. The dashed boxes indicate the adjacent ±1 base positions relative to each breakpoint. Separate models were trained for each of three breakpoint classes. (**b**) MuRaL-indel network architecture. MuRaL-indel employs a U-Net architecture based on convolutional neural networks. The input is the extended sequence context surrounding a breakpoint, one-hot encoded, with a label indicating the occurrence of INDELs of specific lengths at that site. The network’s downsampling path uses five stages of increasing dimensionality to transform the input sequence into a compact, high-dimensional feature map, with convolutional layers extracting features at each step. The upsampling path then gradually restores the spatial resolution via transposed convolutions, combining features from corresponding levels in the downsampling layers through skip connections. The final output is passed through convolutional and fully connected layers to produce a prediction vector of length *n*+1, where *n* is the number of INDEL lengths modeled. The first element (prob[0]) represents the probability of no INDEL, and subsequent elements represent the probabilities of 1-bp, 2-bp, …, up to *n*-bp INDELs at the breakpoint. (**c**) Illustration of two key evaluation metrics: *k*-mer and regional correlations between predicted and observed mutation rates. The top panel shows nucleotide-level predicted rates and observed mutations (e.g., rare variants). The bottom left panel demonstrates 2-mer correlation for 1-bp insertions: observed and predicted rates (obs. *Kᵢ* and pred. *Kᵢ*) are computed for each 2-mer (e.g., ‘AT’, highlighted in the top panel), and their Pearson correlation is calculated across all 2-mers. The bottom right panel shows regional correlation for 100 Kb bins: observed and predicted rates (obs. *Rᵢ* and pred. *Rᵢ*) are averaged per bin, and Pearson correlation is computed across bins. Further details are provided in the Methods.

Unlike MuRaL^25^, which used a ‘local’ module consisting of fully-connected layers and an ‘expanded’ module based on ResNet, MuRaL-indel employs a U-Net architecture based on convolutional neural networks (CNNs) (**Fig. 1b; Supplementary Fig. 1**). The input consists of long sequence contexts surrounding focal breakpoints, one-hot encoded and fed into the network. During training, each breakpoint is labeled based on the presence of an INDEL of a specific length at that site. U-Net’s downsampling, upsampling, and skip connections enable it to capture both local and long-range genomic features influencing INDEL mutagenesis. The final prediction is generated through convolutional and fully connected layers.

The predicted output is a one-dimensional vector of length n+1, where n denotes the predefined number of INDEL length categories modeled (**Fig. 1b**). The first value (‘prob0’) represents the probability of no INDEL mutation occurring at the breakpoint, while subsequent values correspond to the probabilities of INDELs of lengths 1 bp, 2 bp, …, n bp, respectively. Similar to the approach in MuRaL, we applied Dirichlet calibration method^26^ to obtain calibrated probabilities. Furthermore, we employed Poisson distribution correction to address biases arising from possible mutation recurrence when the ratio of positive/negative samples is high in the training data (see Methods).

We trained separate models for insertion, deletion start, and deletion end breakpoints, referred to as the insertion model, deletion start model, and deletion end model, respectively. We focused exclusively on mutation probabilities of small INDELs (<=20bp) in autosomes. For analyses requiring aggregation across multiple models, predicted rates were scaled to match the *de novo* INDEL mutation spectrum prior to aggregation (Methods).

During training and model selection, we evaluated performance using the cross-entropy loss on validation data. Following MuRaL’s evaluation strategy, we further assessed model accuracy by computing the Pearson correlation coefficient between observed and predicted mutation rates, averaged over *k*-mers or binned genomic regions (**Fig. 1c**; see Methods for more details). Using the two correlation metrics, we compared the performance of MuRaL-indel models with different configurations, as well as between MuRaL-indel and existing models. Additionally, we evaluated predictive accuracy by comparing the coefficients of variation (CV) of observed and predicted mutation rates across binned genomic regions, where smaller differences in CV indicate better agreement and improved performance.

### Training and evaluation data for building MuRaL-indel models for humans

Given the abundance of published INDEL variants in human populations, we first trained and evaluated MuRaL-indel using human INDEL datasets. Due to the limited availability of human *de novo* INDELs, we used rare population variants as proxies for mutation events. Specifically, we sampled low-frequency INDELs with varying allele frequencies from the large-scale gnomAD dataset^27^ and compared their mutation spectra to those of *de novo* INDELs to assess representativeness (**Supplementary Fig. 2;** Methods). We found that the spectra of lower-frequency INDELs closely resemble that of *de novo* events, supporting the use of the rarest dataset—‘1in100000’ (see Methods)—for model training and evaluation. Given the scarcity of longer INDELs, we grouped them into seven classes: INDELs of 1–6 bp were modeled individually by length, while those of 7–20 bp were pooled into a single class. This stratification balances resolution for short INDELs with the need to mitigate data sparsity for longer events, enabling robust training and meaningful evaluation.

### MuRaL-indel builds effective mutation rate models

Model performance varies considerably across configurations. We systematically evaluated four key parameters in MuRaL-indel: input sequence length, first CNN layer channel size, training set size, and positive-to-negative sample ratio. Based on validation loss, training efficiency, and *k*-mer/regional correlations, we selected top-performing configurations for downstream analyses (**Supplementary Fig. 3**). Unless otherwise noted, models used an 8 Kb input (4 Kb flanking each side) and 8 channels in the first CNN layer (**Supplementary Fig. 4-5**). For deletion start and end models, we used 4 million samples (2 million deletions, 2 million non-mutated sites; **Supplementary Fig. 6**). The insertion model used a larger dataset of 11 million samples (1 million insertions, 10 million non-mutated sites), as increasing negative sampling improved performance (**Supplementary Fig. 7**). To avoid the influence of strong selective pressure in exon regions, we trained models with only non-exonic rare INDELs (**Supplementary Fig. 8**).

We evaluated model performance for each mutation class using both *k*-mer and regional correlations. We found that most mutation types with varying lengths in the insertion and deletion models exhibited relatively high correlations (**Fig. 2a; Supplementary Fig. 9**). In terms of *k*-mer correlations, as the *k* value increased, the overall correlations slightly decreased, due to fewer observed INDEL mutations in longer *k*-mers. A similar trend is observed in the regional correlation analysis, where varying window sizes also affect the correlation results. Compared to the insertion model, the deletion start and end models demonstrated higher stability in prediction performance across *k*-mers.

**Fig. 2.**
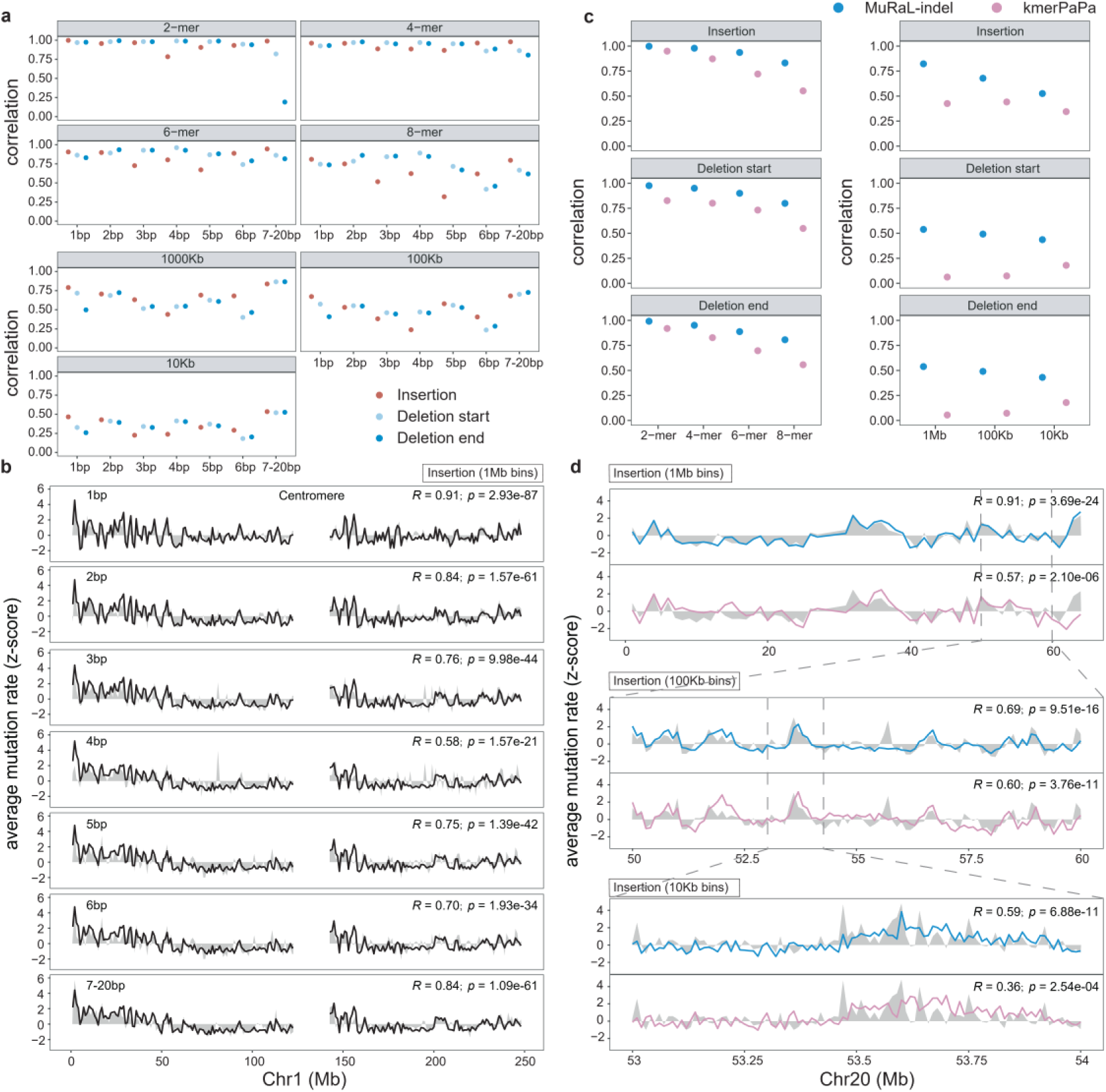
Performance evaluation of MuRaL-indel and existing models. **(a)** *K*-mer and regional correlations for different INDEL types, based on autosomal base-resolution mutation rates predicted by MuRaL-indel. Observed rates were derived from ‘1in100000’ rare variants. **(b)** Example of regional mutation rate correlations at 1 Mb resolution on Chr1, showing observed (gray shading) and predicted (black lines) rates across INDEL types. Rates were z-score normalized for visualization due to differing magnitudes. Centromeric regions lack data. Pearson correlation coefficients and *P* values (two-sided test, no multiple testing correction) are shown in the top-right corners. **(c)** Comparison of *k*-mer and regional correlations between MuRaL-indel and kmerPaPa, using the same observed dataset (‘1in100000’ rare variants). **(d)** Regional correlations at multiple scales on Chr20, with observed rates (gray) and model predictions (colored lines). Color scheme for the lines is the same as panel c. Z-score normalization was applied for comparability; centromeric regions are excluded. Correlation coefficients and *P* values (two-sided) are indicated in the top-right corners.

Regional correlation analyses showed that all three models attained Pearson coefficients around or above 0.5 at 1 Mb resolution, with values nearing 0.5 at finer scales (100 Kb and 10 Kb) (**Fig. 2a**). Beyond genome-wide correlation analysis, we further visualized the mutation rate predictions of different mutation classes within the same genomic regions (**Fig. 2b**). Notably, MuRaL-indel effectively captures local variation in mutation rates, demonstrating high predictive performance across INDELs of diverse lengths.

### MuRaL-indel outperforms existing models

We further compared MuRaL-indel with the recently published kmerPaPa method, which used short sequence contexts (8 bp) to estimate local INDEL mutation rates for the human genome. MuRaL-indel achieved significantly higher correlations than kmerPaPa, highlighting its superior performance (**Fig. 2c; Supplementary Fig. 10**). Particularly at larger genomic scales, MuRaL-indel exhibited markedly improved predictive accuracy. Notably, for the models targeting deletion start and end breakpoints, kmerPaPa exhibited regional correlations close to zero at both 1 Mb and 100 Kb window sizes, suggesting that short-sequence features have inherent limitations in capturing mutation rate landscapes at larger genomic scales (**Fig. 2c**). By comparing mutation rate predictions from both models within the same genomic regions, we found that MuRaL-indel has a more sensitive ability to capture regional mutation rate variation (**Fig. 2d**). Even at a small window size of 10 Kb, MuRaL-indel still maintains a correlation of approximately 0.5.

On the other hand, we assessed the discrepancy between model-predicted and observed regional mutation rates by comparing their coefficients of variation (CV). We found that, across different mutation lengths, the CVs of predicted regional mutation rates in MuRaL-indel were generally close to those observed in real data at both 1 Mb and 100 Kb scales (**Supplementary Fig. 11**). Although some noticeable differences remained at the finer 10 Kb scale, this may be partially attributed to sampling noise due to the sparsity of INDEL mutations in smaller genomic intervals. Nevertheless, the substantial divergence between the CVs of observed and predicted mutation rates points to an area for future improvement.

We further compared the deviation in CV between MuRaL-indel and kmerPaPa relative to the observed values. Our results show that, for the insertion model, MuRaL-indel achieved better agreement with the observed CVs (**Supplementary Fig. 12**). In contrast, both MuRaL-indel and kmerPaPa exhibit considerable room for improvement in capturing the variation in deletion start and end sites.

### Generating mutation rate maps for other species

To evaluate cross-species generalizability, we used MuRaL-indel to predict INDEL mutation rates in three non-human species (*Macaca mulatta, Drosophila melanogaster* and *Arabidopsis thaliana*).

The rhesus macaque (*M. mulatta*) is a close evolutionary relative of humans and a widely used model organism. Given the high genomic similarity and conserved mutational processes between the two species, we reasoned that transfer learning would be effective in this context. Using rare INDELs from a dataset of 572 individuals^28^, we trained both *ab initio* and transfer learning MuRaL-indel models for rhesus macaque (**Supplementary Table 3;** Methods). Transfer learning models initialized their weights from pre-trained human models and used only 40% of the training data required for *ab initio* models. Despite reduced data and shorter training time, transfer learning models achieved comparable performance, with faster and more stable convergence in several cases (**Supplementary Fig. 13**). Notably, they outperformed *ab initio* models at deletion start and end sites (**Fig. 3a-b; Supplementary Fig. 13-14).** These results demonstrate that transfer learning is a more efficient and effective approach when a high-quality pre-trained model exists for a closely related species.

**Fig. 3.**
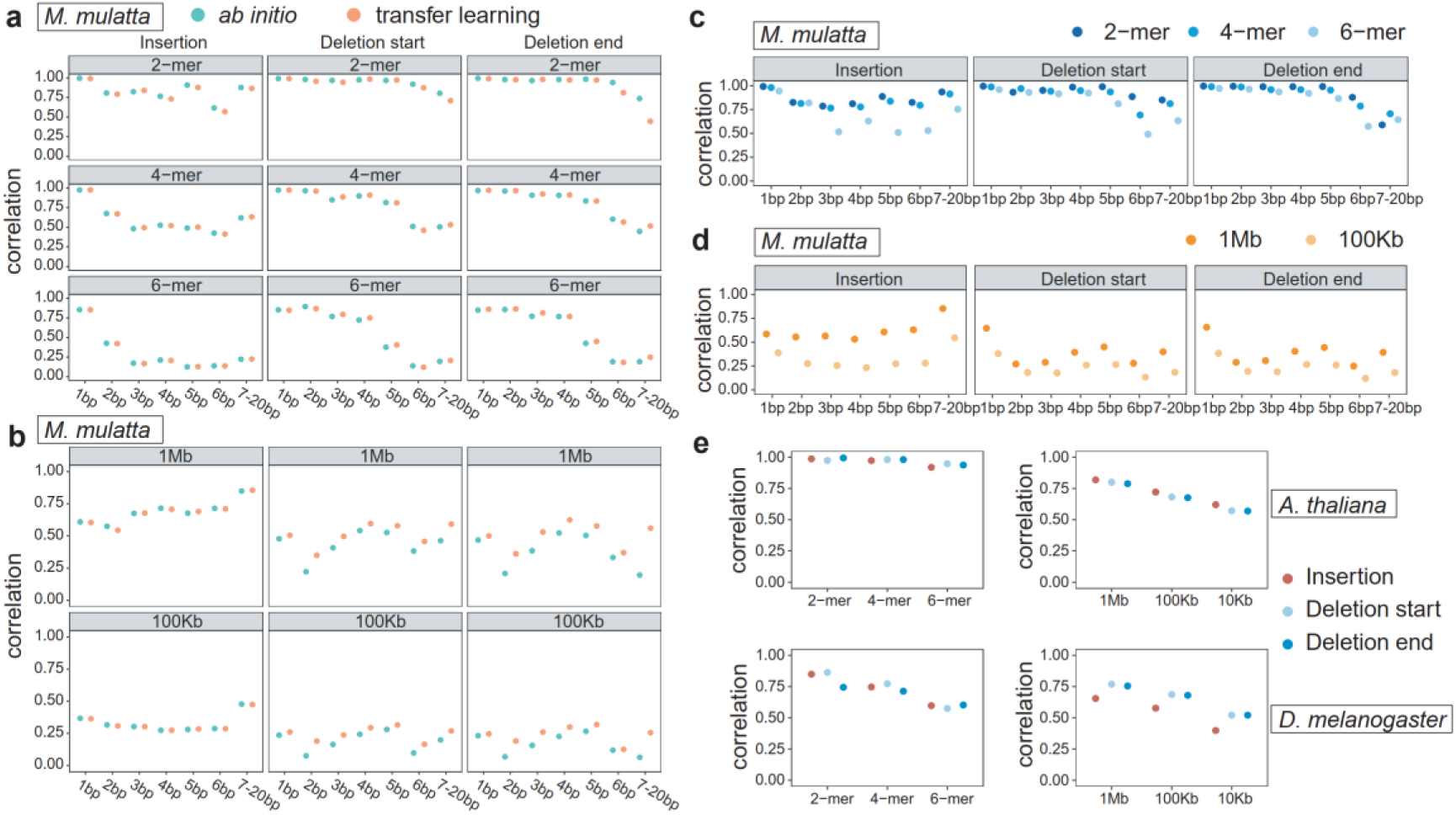
Generating mutation rate maps for other species. (**a**) Evaluation by k-mer correlations for two kinds of models for *M. mulatta*: *ab initio* and transfer learning models. 2-, 4- and 6-mer mutation rate correlations for different mutation types, based on predicted base-resolution mutation rates on rheMac10Plus Chr20. Rare variants of *M. mulatta* were used for calculating observed mutation rates. (**b**) Regional mutation rate correlations with bin sizes of 1 Mb and 100 Kb on rheMac10Plus Chr20 for different mutation types. (**c**) 2-, 4-, and 6-mer mutation rate correlations for different mutation types, based on predicted base-resolution mutation rates on rheMac10Plus whole genome for the *M. mulatta* transfer learning models. (**d**) Regional mutation rate correlations with bin sizes of 1 Mb and 100 Kb for the *M. mulatta* genome. (**e**) *K*-mer and regional correlations for *A. thaliana* and *D. melanogaster*, based on predicted base-resolution mutation rates of the whole genome.

In addition, we trained MuRaL-indel models for two model organisms that are more evolutionarily distant from humans: *D. melanogaster* and *A. thaliana*. Due to their much smaller genome sizes (less than 200 Mb) and relatively small numbers of observed INDELs, we did not further stratify INDELs by length, as was done in humans. Instead, we built models for overall INDEL mutation rates in each species. For both *A. thaliana* and *D. melanogaster*, we trained each model using no more than 50,000 mutations (**Supplementary Table 4-5;** Methods). Despite the relatively small training datasets, the resulting models achieved strong predictive performance, demonstrating that MuRaL-indel is effective even in species with limited mutation data (**Fig. 3e**). At a window size of 10 Kb, regional correlations for most mutation classes were above 0.5, indicating robust applicability of our method in these organisms.

### The distribution of INDEL mutation rates around human genes

We applied the human INDEL mutation rate map to downstream analyses, beginning with an examination of mutation rate profiles across coding regions and their flanks. Based on the variant density of all INDELs and low-frequency INDELs (‘1in100000’) from gnomAD, we generated meta-gene variant density profiles across coding and surrounding regions (Methods). We also plotted INDEL mutation rate profiles around human coding genes using MuRaL-indel predictions (**Fig. 4a**). For most INDEL length types, the observed INDEL density within coding regions was lower than that in both upstream and downstream regions—even in the 1in100000 low-frequency dataset (**Fig. 4a; Supplementary Figs. 15-17**). This pattern was particularly pronounced for non-triplet multiples, which are more likely to cause frameshift mutations. Given the deleterious nature of many coding INDELs, the reduced numbers of INDELs in these regions likely reflect long-term natural selection favoring sequences with lower intrinsic mutability. Overall, the predicted mutation rate profiles for non-triplet INDELs closely mirrored the observed variant density patterns (**Fig. 4a; Supplementary Figs. 15-17**). However, for certain mutation types—such as 3-bp INDELs—a discrepancy between predicted and observed rates remains, highlighting an area for future model refinement.

**Fig. 4.**
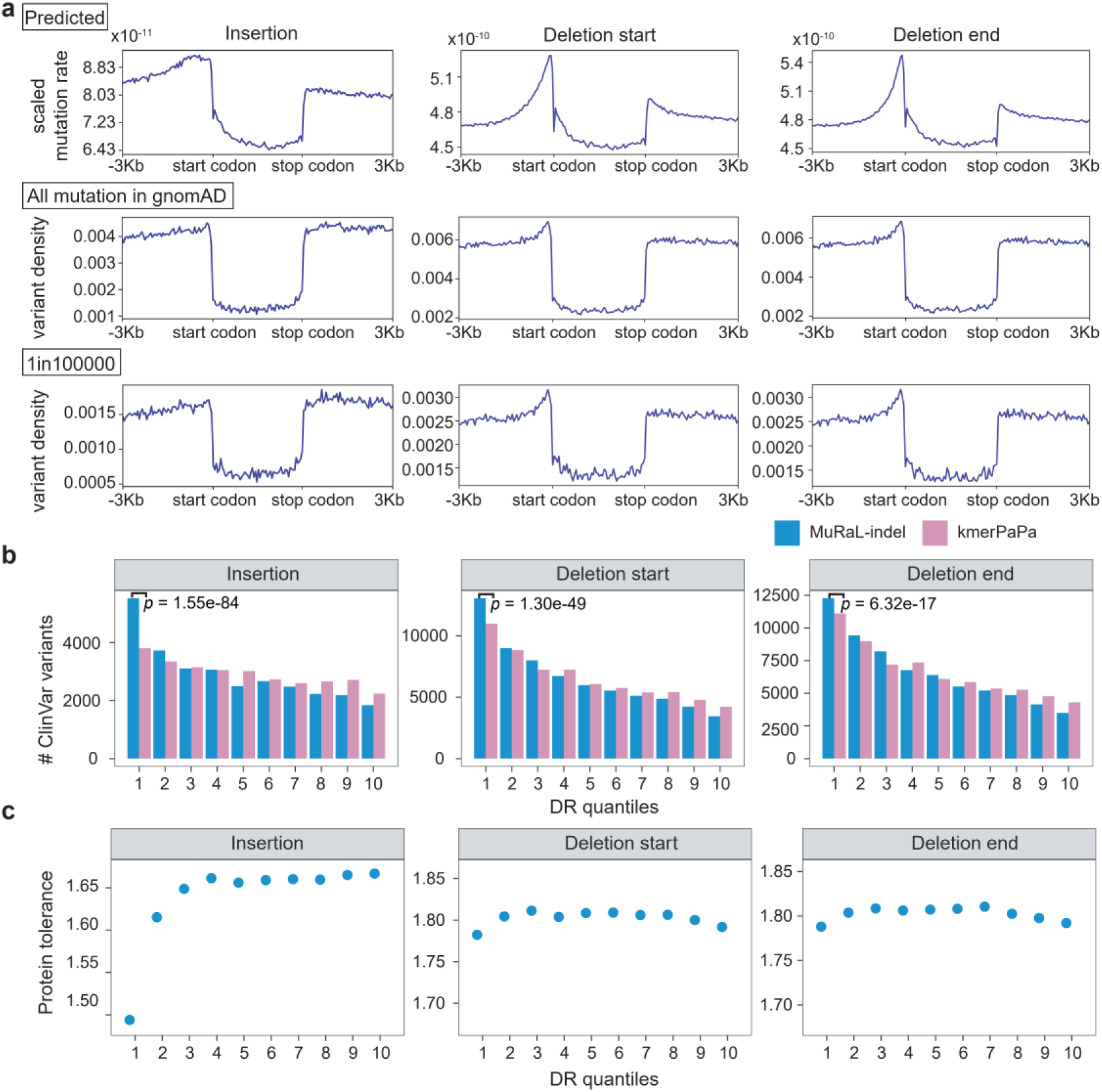
Meta-gene mutation rate profiles around human coding genes and INDEL tolerance analysis. (**a**) Meta-gene mutation rate plots with a bin size of 50 bp for regions around human coding genes, with gene bodies uniformly scaled to 3 Kb. The left was based on scaled MuRaL-indel mutation rates for 1-bp INDELs (see Methods); the center was based on all observed 1-bp INDELs in gnomAD; and the right was based on ‘1in100000’ rare 1-bp INDELs. (**b**) Numbers of ClinVar variants (’Pathogenic’ and ’Likely pathogenic’ in annotations, n = 104,446 ClinVar variants used for analysis) in regions of different DR scores. DR scores were calculated using MuRaL-indel and kmerPaPa mutation rates, respectively. Two-sided Fisher’s exact test was used for comparing the numbers of variants in regions with lowest 10% DR scores. (**c**) Average protein tolerance scores of one-amino-acid INDELs in regions of different DR scores. DR scores were calculated using MuRaL-indel mutation rates.

### Inferring the INDEL tolerance landscape in the human genome

Another downstream application of the predicted mutation rates is to infer the genome-wide INDEL tolerance landscape, which can help prioritize disease-associated variants. Using MuRaL-indel-predicted mutation rates and observed INDELs from gnomAD, we applied the depletion rank (DR) method—previously described in the UK Biobank study^29^—to compute mutational tolerance scores across sliding 500-bp windows in the human genome (**Methods**). We found that regions with the lowest DR scores were significantly enriched for pathogenic INDEL variants from ClinVar (**Fig. 4b**). Notably, this enrichment was substantially stronger when DR scores were computed using MuRaL-indel mutation rates compared to those with kmerPaPa, indicating that MuRaL-indel–based DR scores provide informative signals for prioritizing disease-associated variants.

Furthermore, we analyzed the data of a recently proposed single-amino-acid INDEL tolerance scoring model^30^, which estimates tolerance scores for insertions and deletions at every amino acid position across all human-expressed proteins. DR scores based on 3-bp INDEL mutation rates by MuRaL-indel revealed that the lowest-ranked bins were enriched for more conserved amino acid sites—a trend particularly evident in insertions (**Fig. 4c**; Methods). In contrast, deletions exhibited generally higher tolerance levels, suggesting single-amino-acid deletions tend to be less deleterious than single-amino-acid insertions.

### Sequence motifs associated with high INDEL mutation rates

To gain insight into the mutational mechanisms, we next investigated sequence features associated with high INDEL mutability in the human genome based on predictions of MuRaL-indel (see Methods). We used Integrated Gradients (IG)^31^, a method designed specifically for deep learning models, to compute contribution scores of input sequence features. Due to computational constraints, we focused on 1-bp and 2-bp INDELs, analyzing the top 100,000 genomic positions with the highest predicted mutation rates (designated hypermutated sites), with 100,000 randomly sampled positions as controls. IG analysis revealed that high-contribution regions within the central 200 bp captured most of the predictive signal in the 8 Kb input context (**Fig. 5a** and **Supplementary Fig. 18**, see Methods for contribution calculation). We further used MoDISco^32^ to identify motifs associated with high contribution scores within the central 200 bp of hypermutated sites, termed seqlets (**Supplementary Figs. 19-21**). Genomic regions containing these seqlets exhibit elevated rare variant densities compared to most 8-mers, supporting their designation as genuine mutation hotspots (**Fig. 5b**, **Supplementary Fig. 22**).

**Fig. 5.**
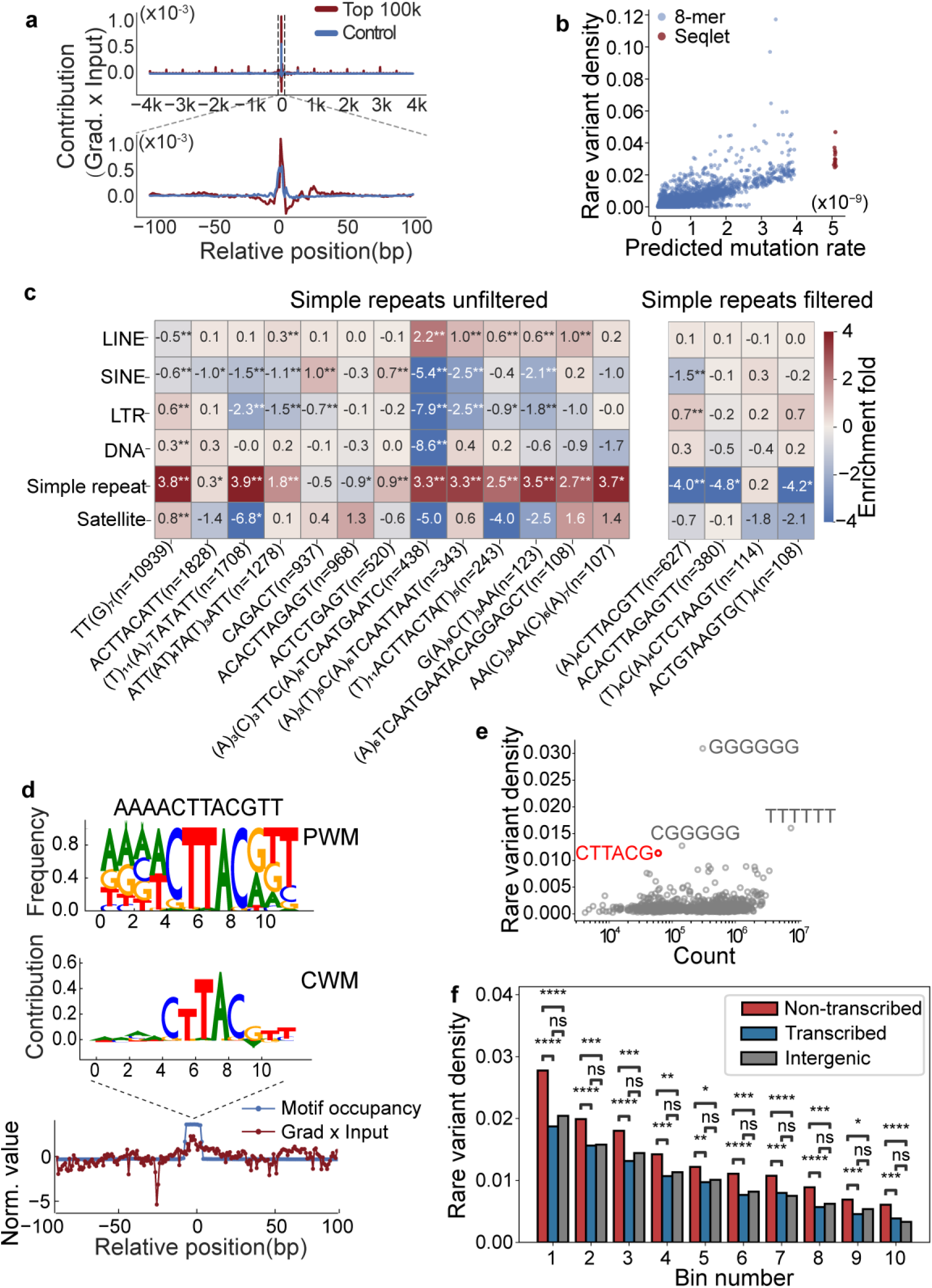
| Motifs associated with 1-bp insertion hypermutability. (**a**) Average base-level contribution (gradient × input) within ±4 Kb (upper) or ±100 bp (lower) of focal sites. red: hypermutated sites; blue: control sites. (**b**) Validating the hypermutability of identified motifs (seqlets) by rare variant densities. Red dots represent average rare variant densities of hypermutated sites whose ±100 bp flanks contain a specific seqlet; blue dots represent average rare variant densities of specific 8-mer motifs. (**c**) Repeat enrichment analysis of motifs, with the left panel displaying results before filtering and the right panel showing results after excluding hypermutated sites overlapping simple repeats. The motif counts are given in the brackets. Motifs in the form of (X)ₙ indicate n consecutive occurrences of base X, e.g., (T)₁₁ represents 11 consecutive thymine bases. (**d**) Position weight matrix (PWM), motif contribution weight matrix (CWM), and motif occurrence, mean contribution across the central 200 bp, after excluding simple repeats. The PWM was generated with all found motif instances in the top 100,000 sites. The CWM was obtained by multiplying the seqlets with their corresponding gradients and averaging across seqlets. (**e**) Genome-wide rare variant densities versus 6-mer occurrence counts. Each point represents a unique 6-mer together with its reverse complement. (**f**) Strand-specific rare variant densities for the CTTACG motifs in different bins stratified by predicted mutation rates (first bin with highest mutation rates). ‘**’, p < 0.01; ‘****’, p < 1 × 10⁻⁴ (Fisher’s exact test).

For 1-bp INDELs, motifs associated with hypermutated sites are mainly mononucleotide repeats and AT-rich elements, whereas motifs for 2-bp INDELs were enriched for dinucleotide repeats (**Supplementary Figs. 19-21**). These motifs were predominantly found in simple repeat regions (**Fig. 5c; Supplementary Fig. 23**), suggesting that polymerase slippage during replication is likely the major mechanism causing high mutation rates at the sites. To identify non-repeat motifs, we repeated the enrichment analysis excluding hypermutated sites overlapping simple repeats and still detected several enriched motifs for 1-bp insertions and 2-bp deletion starts (**Fig. 5c**; **Supplementary Fig. 24**). We focused on the 1-bp insertion motif—AAAACTTACGTT—which had the highest motif count (n = 627) and was significantly enriched in long terminal repeats (LTRs) (**Fig. 5c**). The position weight matrix (PWM) of this motif indicates little repeat content (**Fig. 5d**), suggesting that replicative slippage is unlikely to drive elevated mutation rates in these regions. Occurrences of this motif were enriched in the central regions of hypermutated sites and correspondingly exhibited high contribution scores in those regions (**Fig. 5d**), implying that the motif is the primary cause for the elevated mutability.

We further investigated CTTACG (or its reverse complement CGTAAG), the core hexamer of this motif identified from the contribution weight matrix (CWM) (**Fig. 5d**). The observed genome-wide rare variant density of CTTACG ranked fourth among all 2,048 possible 6-bp motifs for 1-bp insertions (**Fig. 5e**) and predominantly mutated to CTTAACG (**Supplementary Fig. 25**). A similar motif (‘GTAAGT’) was previously reported in early studies of INDEL hotspots^33,34^, yet the underlying mutational mechanism has remained unknown. When divided the CTTACG sites of the human genome into 10 bins based on their predicted mutation rates, the top 8 bins were significantly enriched in promoters, UTRs, and exons. Intriguingly, each bin showed elevated rare variant density preferentially on the non-transcribed strand, suggesting a link to transcription-associated DNA damage or repair (**Fig. 5f**; **Supplementary Fig. 25**). Furthermore, the reverse complement of CTTACG closely matched the consensus 5′ splice site (5SS) consensus sequence (GTAAG)^35^, and highly mutable CTTACG sites were enriched for 5SS locations, implicating that co-transcriptional splicing may contribute to increased mutability of this motif (**Supplementary Fig. 25**).

We also found that the CTTACG motif is also associated with high 1-bp deletion rate in humans (**Supplementary Fig. 26**). Furthermore, elevated INDEL rare variant densities in the CTTACG context were also observed in rhesus macaque, but not in Drosophila or Arabidopsis **(Supplementary Fig. 26**), suggesting that the hypermutability of CTTACG motifs may be specific to mammalian or primate lineages.

## Discussion

Small insertions and deletions (INDELs) are critical sources of genetic diversity and disease. Accurate estimation of their mutation rates has been hindered by data scarcity, subtype diversity, and the inability of existing models to capture long-range genomic context. To address this, we developed MuRaL-indel, a deep learning framework that predicts germline INDEL mutation rates by leveraging long-sequence contexts (up to 8 Kb) via a U-Net architecture. Using rare variants from gnomAD as a proxy for *de novo* mutations, we generated high-resolution, length-stratified INDEL mutation rate maps for the human genome. Comprehensive benchmarking demonstrated that MuRaL-indel outperforms existing methods like kmerPaPa across scales from kilobases to megabases. The model’s generalizability was validated across multiple species. For rhesus macaque, transfer learning from the human model proved more efficient than *ab initio* training. We also successfully applied MuRaL-indel to more distant species (*D. melanogaster*and *A. thaliana*), achieving high predictive performance even with limited data. Downstream applications confirmed the utility of the predicted maps. MuRaL-indel recapitulated the selective constraint on coding regions, particularly for frameshift-inducing non-triplet INDELs. Furthermore, mutation tolerance scores derived from its predictions showed stronger enrichment for pathogenic ClinVar variants in constrained regions than kmerPaPa, enhancing disease variant prioritization.

Mechanistic insights were gained by identifying sequence motifs linked to hypermutability. While mononucleotide and dinucleotide repeats dominated 1-bp and 2-bp INDEL hotspots—consistent with polymerase slippage—we also discovered several non-repeat motifs. Particularly, the non-repeat motif containing a core hexamer CTTACG associated with 1-bp insertion hypermutability exhibits strand asymmetry and enrichment in genic regions, suggesting a link to transcription-associated mutagenic processes. Its conservation in rhesus macaque but not in *Drosophila* or *Arabidopsis* indicates a potentially mammal-specific mutagenic mechanism.

Despite these advances, several limitations remain to be addressed. Prediction accuracy declines in repetitive or low-mappability regions such as centromeres and telomeres, where available data and sequence features are limited. INDELs longer than 6 bp remain underrepresented and are modeled only coarsely; more data or tailored architectures could improve this range. Currently, MuRaL-indel relies exclusively on sequence information, which supports generalization but omits other relevant genomic features such as chromatin state, replication timing, or transcriptional activity. Integrating such data could capture residual variance in mutation rates. Finally, extending the model to additional species requires careful handling of assembly and variant-calling artifacts that might distort estimates.

Several promising directions emerge from this work. Future research could enhance the model by incorporating non-sequence features like chromatin accessibility, replication timing, or transcriptional levels. These contextual factors are essential for refining predictions, especially in complex genomic regions. Exploring advanced deep learning methods may also improve the capture of long-range mutability dependencies. Biologically, future studies may clarify the precise molecular mechanisms driving the hypermutability of the non-repeat CTTACT motif through targeted *in vitro* or cellular assays. It will also be important to investigate other non-repeat motifs associated with hypermutability of other INDEL subtypes. Additionally, extending the framework to model somatic INDEL mutations in cancer could contribute to translational and precision medicine applications.

In summary, MuRaL-indel provides a generalizable framework for high-resolution INDEL mutation rate mapping. Its superior performance and practical utility in evolutionary and biomedical analyses make it a valuable resource for the community, poised to advance INDEL mutation biology and genomic medicine.

## Methods

### Design of the MuRaL-indel model

Previous studies have shown that INDEL mutation rates are influenced by both local nucleotide context and broader sequence features. Additionally, longer INDELs are less frequently observed and often omitted from predictive models. To learn INDEL mutability signals from the sequence context of a focal site, a neural network with a U-Net architecture was constructed to learn signals at multiple scales through upsampling and downsampling processes. In the downsampling process, the model has five sequential dimensionality-expansion stages, transforming a low-dimensional, large-scale matrix of multi-kilobase sequences into a high-dimensional, compact feature representation. At each stage, convolutional layers extract spatial features following dimensionality expansion, followed by residual blocks to preserve gradient flow and enhance learning stability. The residual blocks are based on the ResNet architecture^36^. The upsampling process restores the high-dimensional compact representation to the original sequence resolution through upsampling operations. During this process, convolutional layers and residual blocks refine the features, while skip connections integrate spatially matched intermediate states from the downsampling process. Final predictions are generated via additional convolutional and fully connected layers, outputting a (n+1)-dimensional vector for each site. Here, n represents the predefined number of INDEL length categories. The vector’s first element denotes the probability of no INDEL occurrence, while subsequent elements correspond to the probabilities of 1-bp, 2-bp, …, n-bp INDELs at the junction (or breakpoint) between adjacent nucleotides.

The key architectural hyperparameters of MuRaL-indel include: input sequence radius, downsampling stride list, feature channel numbers and convolutional kernel size. First, the input sequence length is determined by the radius, which specifies the number of nucleotides flanking each side of the central site. The downsampling stride list controls the spatial compression in the U-Net encoder. The first value is fixed at 1, while the remaining five are adjusted based on the input radius to ensure effective multi-scale feature extraction across different resolution levels. This same list is used at the later upsampling stage, enabling symmetric feature reconstruction via skip connections. The channel number parameter sets the initial number of feature maps, with subsequent layers increasing the channel number linearly through dimension-expansion stages. Finally, the kernel size is selected according to the input radius and downsampling strides to balance model capacity and computational efficiency, ensuring robust local pattern learning without excessive parameter overhead. (see **Supplementary Fig. 1**).

### Model implementation

We implemented the MuRaL model with PyTorch framework^37^, along with APIs from pybedtools^38^. For model training, we used the cross-entropy loss function and the Adam optimizer^39^ for learning model parameters, and employed Ray Tune^40^ to facilitate hyperparameter tuning (**Supplementary Fig. 1**). The scheduler ‘ASHAScheduler’ in Ray Tune was used to coordinate trials and execute early stopping before reaching the specified maximum number of training epochs (e.g., 10), which can substantially reduce the training time. The mean cross-entropy loss of the validation sites (termed validation loss) was calculated at the end of each training epoch. We further set a stopping rule to terminate a trial if three consecutive epochs did not obtain a validation loss smaller than the current minimum validation loss. The ‘learning rate’ and ‘weight decay’ of Adam optimizer were two hyperparameters that could affect the learning performance significantly. We used Ray Tune to run trials with different values for the two hyperparameters. To have better convergence, we used the learning rate scheduler ‘lr_scheduler.StepLR’ in PyTorch to decay the learning rate after a number of steps by a specified factor.

### Human mutation data for model training and evaluation

#### Rare variants from gnomAD

Rare variants typically arose more recently in evolutionary time and are subject to weaker influence from natural selection and nonadaptive evolutionary forces compared to common variants. Previous studies^19,41,42^ have demonstrated that rare single-nucleotide variants (SNVs) can serve as reliable proxies for mutation rate estimation. Like the MuRaL^25^ framework for modeling point mutations, we applied a downsampling strategy to polymorphism data to mitigate potential mutational hotspot saturation, which is common in large-scale datasets where high-frequency INDELs may distort rate estimates due to recurrent mutations or sequencing artifacts. Specifically, we focused on low-frequency INDELs from the gnomAD database (v3.1.1)^27^, which aggregates whole-genome sequencing data from 76,156 individuals. Only small INDELs in autosomes were considered, as other mutation types and sex chromosomes have specific features that need to be modeled separately. We extracted rare INDELs from gnomAD to approximate *de novo* INDELs.

When the sample size is as large as that of gnomAD, sequence contexts with high mutation rates—such as simple repeats—may approach mutational saturation, increasing the likelihood of recurrent (independent) mutations at the same genomic position. To mitigate this, we downsampled the gnomAD data to a specified total allele count using a hypergeometric distribution, and randomly generated the corresponding alternative allele counts from the same distribution (see probability density function below):

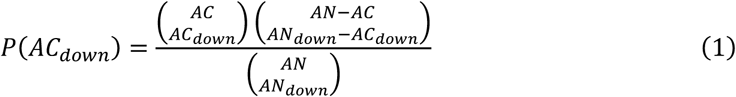

Here, AN and AC denote the total and alternative allele counts in the original data, respectively, while AN_down_ and AC_down_ represent the corresponding counts after downsampling. For each polymorphic site, given AN, AC, and AN_down_, we sampled AC_down_ from a hypergeometric distribution as defined in Equation (1). Variants were then filtered to retain those with a specific alternative allele frequency (i.e., AC_down_ / AN_down_) in the downsampled data, forming the rare variant datasets used in subsequent analyses.

First, we downsampled the gnomAD data to total allele counts of 200, 2,000, 20,000, and 100,000 (corresponding to 100, 1,000, 10,000, and 50,000 diploid genomes, respectively) and extracted singleton INDELs from each dataset (corresponding to allele frequencies of 1/200, 1/2,000, 1/20,000, and 1/100,000). These four rare INDEL datasets were designated ‘1in200’, ‘1in2000’, ‘1in20000’, and ‘1in100000’, respectively. In cases where multiple alternative alleles at the same position met the rare variant criterion in a downsampled dataset, only one was randomly selected for downstream analyses to ensure no position contained more than one rare variant. The number of rare INDELs in each dataset is summarized in **Supplementary Table 1**.

#### De novo mutations

We collected *de novo* INDELs from the gene4denovo database (v1.2)^43^ for analysis. Because some data sources in gene4denovo database contributed only a small number of *de novo* INDELs and different studies used distinct methods for variant calling, we used only the *de novo* INDELs from three large-scale studies^44–46^ for our analysis, which consisted of 46,623 unique *de novo* INDELs.

To assess whether the extracted rare INDELs accurately reflect the properties of de novo INDELs, we compared their mutation spectra. We counted the occurrences of different INDEL lengths and 2-mer mutation types for each dataset and calculated the relative proportion of deletions and insertions in the specific dataset. We found that mutation spectra of rare INDELs were highly similar with that of *de novo* INDELs (**Supplementary Fig. 2**). When the downsampling frequency decreases, rare INDELs increasingly resemble *de novo* INDELs in both length distribution and 2-mer sequence context composition (**Supplementary Fig. 2**), so we chose the rarest INDEL dataset (‘1in100000’) for model training and evaluation.

In the original INDEL data, repetitive sequences can introduce ambiguity in pinpointing the exact insertion or deletion locations. We referred to the approach proposed by Bethune et a_l_^21^ to resolve this, applying an enumeration-based realignment strategy and randomly selected one plausible position as the final assigned site.

Genomic regions with excessively low or high read coverage are prone to false positive and false negative variant calls. To mitigate this, we used gnomAD coverage data to exclude positions outside the optimal range. The genome-wide mean coverage per individual in gnomAD is 30.5×, and we retained autosomal positions with coverage between 15× and 45×—totaling 2,626,258,019 base pairs—as high-mappability sites for downstream analyses.

#### Training and validation data for human MuRaL-indel models

For the human dataset, we trained separate models for insertion sites, deletion start sites, and deletion end sites. Specifically, we randomly selected 1,000,000 rare insertion mutations and 10,000,000 non-mutated sites to train the MuRaL-indel insertion model. For the deletion models, we used 2,000,000 rare deletion mutations and 2,000,000 non-mutated sites to construct the training sets for both deletion start and end sites. During training, an independent validation dataset, comprising 10% of the size of the corresponding training set, was used to evaluate model performance. The configuration of key hyperparameters for human MuRaL-indel models was provided in **Supplementary Table 7**.

### Calibrating and scaling predicted probabilities

As demonstrated in the MuRaL framework^25^ for point mutations, Dirichlet calibration effectively reduces miscalibration errors in multi-class classification tasks. This method has been shown to improve performance across key metrics, including expected calibration error (ECE), classwise-ECE, and Brier score^26^. We adopt the same Dirichlet calibration procedure to refine the raw probability estimates from our U-Net model, ensuring that the predicted INDEL length distributions align with the true empirical frequencies.

Due to the sparse genomic distribution of INDELs, training on datasets reflecting the true INDEL density (#INDELs / #all genomic sites) would result in highly imbalanced data. This would not only lead to sparse feature representations but also incur substantial computational costs. To address this, we employed a class-balanced training strategy where positive (INDEL) and negative (non-INDEL) samples were subsampled at relatively balanced ratios. However, this artificial class imbalance in training data may bias predicted probabilities toward overestimating rare INDEL events or underestimating common ones. To correct for this bias, we further applied a Poisson-based calibration. The Poisson calibration is derived as follows:

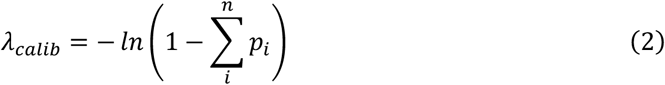

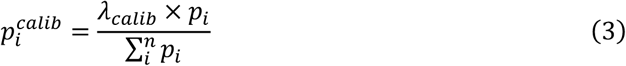

Here, n is the number of mutation subtypes and p_i_ is the estimated probability of a specific subtype.

The absolute values of above mutational probabilities were not mutation rates per bp per generation. To obtain a mutation rate per bp per generation for each nucleotide, we may scale the calibrated probabilities using previously reported genome-wide *de novo* INDEL mutation rate and spectrum per generation. The scaling factor (𝑓^𝑚𝑢𝑡^in the equations below) can be calculated using following equations:

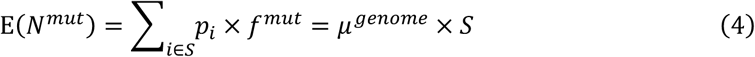

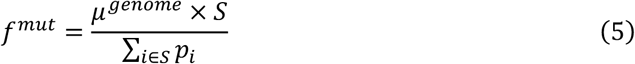

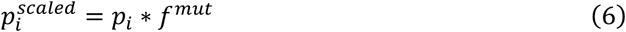

where 𝐸(𝑁^𝑚𝑢𝑡^) is the expected number of INDEL mutation, 𝜇^𝑔𝑒𝑛𝑜𝑚𝑒^is the per-base per-generation mutation rate obtained from *de novo* INDEL mutation data, |𝑆| is the number of considered genomic sites (𝑆) for calculating scaling factors. 𝑝_𝑖_ and 𝑝^𝑠𝑐𝑎𝑙𝑒𝑑^are base-resolution mutation probabilities before and after scaling. For the human genome, we used the 𝜇^𝑔𝑒𝑛𝑜𝑚𝑒^ of 1.5 x 10^-9^ (according to a recent study^47^) and validation sites of MuRaL-indel models (𝑆) to calculate the scaling factors, and then generated scaled mutation rates. We note that applying or omitting this scaling step does not affect the calculation of *k*-mer and regional mutation rate correlations described below.

### Evaluation of model performance

In line with the MuRaL framework^25^, we adopted two complementary metrics—*k*-mer correlation and regional correlation—to evaluate model performance at higher levels of aggregation, as direct assessment of per-nucleotide mutation rate accuracy remains infeasible. These metrics provide a validated approach to quantify the concordance between predicted mutation rates and empirically observed genomic patterns across sequence contexts and genomic regions.

#### Correlation analysis of k-mer mutation rates

The *k*-mer correlation metric evaluates the model’s predictive performance in local sequence contexts. Specifically, mutations are categorized into *k*-mer subtypes based on equidistant flanking sequences (i.e., upstream and downstream nucleotides) at each nucleotide junction. For each mutation type (e.g., 1-bp INDELs), the following steps are performed:

For the *i*th *k*-mer subtype, we calculated the observed mutated rate 𝐾^𝑜𝑏𝑠^ and the predicted mutation rate 𝐾^𝑝𝑟𝑒𝑑^ in the considered subtype:

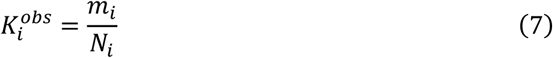

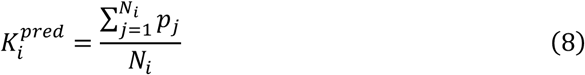

where *m_i_* is the observed number of mutated sites belonging to that *k*-mer subtype, 𝑁_𝑖_ is the total number of sites harboring the reference *k*-mer motif, and 𝑝_𝑗_ is the predicted mutation probability of the *j*th valid site.

Based on the calculated observed and predicted *k*-mer mutation rates, we can calculate the Pearson correlation coefficient for any set of *k*-mer subtypes (one subtype as a datapoint).

#### Correlation analysis of regional mutation rates

While *k*-mer correlation assesses local sequence context dependencies, the regional correlation evaluates the model’s predictive performance across broader genomic domains. This metric quantifies the concordance between predicted and observed mutation rates across non-overlapping genomic bins.

More specifically, for a specific mutation type, we calculated the observed and predicted mutation rates as below.

First, we divided a specified region (e.g., a chromosome) into non-overlapping bins with a given bin size (e.g. 10Kb, 100Kb, etc.). For the *i*th binned region, we calculated the observed mutated rate 𝑅_i_^𝑜𝑏𝑠^ and the predicted mutation rate 𝑅^𝑝𝑟𝑒𝑑^,

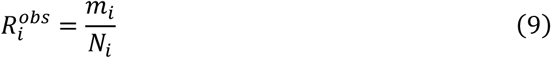

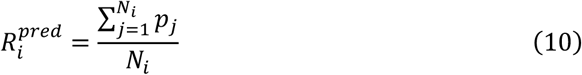

where 𝑚_𝑖_ is the number of observed mutations of the specific mutation type, 𝑁_𝑖_ is the total number of sites with same base as the reference base of that mutation type, and 𝑝_𝑗_ is the predicted mutation probability of the *j*th valid site in the binned region.

Using the calculated observed and predicted regional mutation rates, we computed the Pearson correlation coefficient across binned genomic regions, with each bin treated as a data point. To account for gaps and low-mappability regions, and to avoid unreliable estimates from bins with few valid sites, we restricted the correlation analysis to bins satisfying the criterion 𝑁 > 20% ∗ 𝑁_𝑚𝑒𝑑𝑖𝑎𝑛_, where 𝑁_𝑚𝑒𝑑𝑖𝑎𝑛_ is the median number of valid sites per bin across the chromosome.

### Comparison of models with different hyperparameters

During training of the MuRaL-indel model, several factors—such as input sequence length, model channel dimensions, training set size, positive-to-negative sample ratio, and genomic context—can influence performance. To identify optimal hyperparameters and dataset designs, we conducted systematic ablation studies across these variables.

Input sequence length determines the extent of contextual information available for mutation prediction. While longer sequences may improve accuracy by capturing extended motifs, they also increase computational cost and overfitting risk. To assess this trade-off, we evaluated models with varying input length (1, 2, 4, 6, 8, and 10 Kb) and measured their performance (**Supplementary Fig. 3-4**).

Model channel numbers control representational capacity, particularly in the first convolutional layer of the U-Net backbone; subsequent layers scale proportionally with depth to maintain hierarchical feature extraction (**Supplementary Table 5**). We tested four configurations (8, 16, 24, and 32 channels), comparing predictive performance and parameter count to balance accuracy and computational efficiency (**Supplementary Fig. 3; Supplementary Fig. 5**).

The size of the training dataset is a fundamental determinant of model generalization. While larger datasets typically enhance predictive accuracy by capturing diverse mutation contexts, excessive data may introduce computational burdens, inefficient resource allocation, and overfitting scarce mutation patterns. To identify the optimal trade-off, we evaluated training set sizes ranging from 1 million (M) to 4 M samples in the MuRaL-indel framework (**Supplementary Fig. 3; Supplementary Fig. 6**).

During training of the human insertion model, we found that using the same positive-to-negative sample ratio as the deletion model did not yield optimal performance. To address this, we systematically evaluated the impact of class imbalance by varying the ratio of non-mutated (negative) to insertion (positive) sites (**Supplementary Fig. 3; Supplementary Fig. 7**). We constructed four training datasets with negative-to-positive ratios of 1:1, 2:1, 5:1, and 10:1, corresponding to total sizes of 2 M, 3 M, 6 M, and 11 M samples, respectively. The increasing dataset size ensured adequate coverage of rare insertion sites while proportionally expanding the negative cohort, allowing us to identify the optimal balance between class representation and model performance.

Furthermore, INDELs are subject to stronger selective constraints in coding regions compared to non-coding sequences. Retaining exonic mutation sites in training and validation datasets may introduce selective pressure as a confounding factor, potentially widening the gap between model-predicted mutation rates and true de novo mutation rates. To evaluate how exonic mutation data influences model training, we conducted two parallel experiments: exonic-inclusive training, which retained exonic INDELs in both training and validation datasets, and exonic-exclusive training, which masked exonic regions during data sampling and validation (**Supplementary Fig. 8**).

### Comparison of MuRaL- indel models with previously published models

We compared MuRaL-indel with kmerPaPa^21^, a previously published approach that achieved the highest predictive performance to date. The kmerPaPa model was based on the upstream and downstream 4 bp (8-mer) of INDEL sites. To mitigate data sparsity for longer k-mers, kmerPaPa applied the IUPAC degeneracy rule to group similar motifs. We generated the INDEL mutation rate map of the human genome based on the 8-mer mutation rates inferred by kmerPaPa. Subsequently, we compared the *k*-mer correlation and regional correlation of MuRaL-indel and kmerPaPa models at the whole-genome and on each autosome (**Supplementary Fig. 10**). The kmerPaPa model did not provide length-specific mutation rate predictions, limiting its resolution compared to our approach.

#### Applying MuRaL to other species

We applied MuRaL-indel to predict the INDEL mutation rates of three other species:

1. *M. mulatta, D. melanogaster* and *A. thaliana*.

We obtained INDEL mutation data from a recent *M. mulatta* population study^28^ (1026 individuals), which retained 572 high-quality genomes after rigorous quality control. The raw variant call file (VCF) was publicly accessible at https://rhesusbase.com/download/1kmg/rheMac10Plus_572rhe_remove254,401SNVs.vc f.gz. Given that this study improved the rhesus macaque reference genome ("rheMac10Plus") through long-read sequencing gap-filling, we adopted rheMac10Plus.fa (https://rhesusbase.com/download/1kmg/rheMac10Plus.fa) as the reference for downstream analysis. To ensure high-confidence training data, we re-mapped three deeply sequenced individuals (SRR1791528, SRR1964575, SRR10357711) to rheMac10Plus using bwa-mem2^48^ with default parameters. Coverage depth was calculated by merging BAM files (mean = 113×) and we retained sites within ±50% of the mean (56×–169×). This range balances sensitivity and error suppression in high-depth regions. From the original 9,032,942 INDELs, we applied a downsampling strategy (AN_down_ = 1000) to extract singleton variants (AC_down_ = 1) in autosomes. For multi-allelic site, we randomly selected one variant per locus to avoid overrepresentation. After filtering by coverage depths, we retained 1,004,246 insertion sites, 1,568,420 deletion start sites and 1,561,913 deletion end sites.

We trained *ab initio* models as well as transfer learning models for *M. mulatta*. For each *ab initio* model, we compiled a training dataset consisting of 500,000 mutated and 5,000,000 non-mutated sites, and an independent validation dataset consisting of 50,000 mutated and 500,000 non-mutated sites. We used the same hyperparameter setting as that for human *ab initio* models. For each transfer learning model, we compiled a training dataset consisting of 200,000 mutated and 2,000,000 non-mutated sites and an independent validation dataset consisting of 20,000 mutated and 200,000 non-mutated sites (**Supplementary Table 3**). All models underwent three independent training trials to assess reproducibility, with a maximum of 50 epochs per trial. As transfer learning models performed better than *ab initio* models, we used the trained transfer learning models to generate base-resolution predictions for the whole genome of *M. mulatta*, with *k*-mer correlations and regional correlations shown in **Supplementary Fig. 14**.

We obtained *A. thaliana* INDEL data from the 1001 Genomes Project (https://1001genomes.org/)^49^, which included 12,883,854 polymorphic sites for 1135 inbred lines. Because of long-term inbreeding, the minimum allele count (AC) for polymorphisms is 2, so we retained biallelic INDELs with AC = 2 for analysis. After filtering sequencing depth (mean = 21×) to retain regions within ±50% of the mean (10×–30×), we identified 163,891 insertions and 238,731 deletions, which were insufficient for length-stratified modeling. Consequently, we trained a binary classification model (mutation vs. non-mutation) using 50,000 mutation sites combined with 1,000,000 non-mutation sites (1:20 ratio) for training, and 5,000 mutation sites with 100,000 non-mutation sites for validation (**Supplementary Table 4**).

We obtained *D. melanogaster* INDEL data from the Drosophila Genetic Reference Panel (DGRP)^50^, which includes 3,837,601 polymorphic sites across 205 inbred lines (excluding sex chromosomes and heterochromatic sequences). Following a strategy analogous to *A. thaliana* processing, we retained singleton INDELs for analysis, yielding 23,427 insertions and 56,357 deletions. Due to the low mutation counts, we trained a binary classification model (mutation vs. non-mutation) without length stratification. For insertions, the training set comprised 10,000 mutation sites and 200,000 non-mutation sites (1:20 ratio), with 1,000 mutations and 20,000 non-mutation sites for validation. For deletions, the training set included 20,000 mutation sites and 400,000 non-mutation sites, validated with 2,000 mutations and 40,000 non-mutation sites (**Supplementary Table 5**). The configurations of hyperparameters for MuRaL models of the three species were provided in **Supplementary Table 8-10**. For each training task, the checkpointed model with lowest validation loss among all trials was used for predicting base-wise mutation rates in the whole genome. The calculation of *k*-mer and regional mutation rate correlations was the same as that for the human data.

### Meta-gene profiles of mutation rates around human coding genes

We downloaded human gene annotations from GENCODE^51^ (v44) and extracted the longest transcript for each protein-coding gene as its representative transcript. DeepTools^52^ (v3.5.5) was used for plotting meta-gene profiles and heatmaps for mutation rates around protein-coding regions.

### Depletion rank

We calculated the depletion rank (DR) scores using the same method as that in Halldorsson et al.^29^. For each of sliding 500-bp windows with a 50-bp step across the genome, we computed the observed number of all gnomAD INDELs (O) and the expected number of INDELs (E). The expected number of INDELs (E) was calculated by summing the scaled MuRaL mutation rates in the given window and multiplying the summed rate by a correction factor. The correction factor was the ratio between the observed number of gnomAD INDELs and the summed mutation rate in a set of presumably neutral sites from a previous study^53^. With obtained O and E values, we then calculated the metric (𝑂 − 𝐸)/√𝐸 for all windows. The window with the i-th lowest (𝑂 −𝐸)/√𝐸 was assigned a DR score of 100*(i−0.5)/n, where n is the total number of windows.

The ClinVar INDELs were downloaded from https://ftp.ncbi.nlm.nih.gov/pub/clinvar/vcf_GRCh38/archive_2.0/2025/clinvar_20250209. vcf.gz. We extracted INDELs which were labeled as pathogenic or likely pathogenic. These included 32,107 insertions and 72,339 deletions.

We reanalyzed the data of INDEL^i30^, which quantified INDEL tolerance across protein domains by integrating deep mutational scanning data for nine structurally characterized domains. It provided genome-wide functional impact predictions for single-amino acid INDELs, yielding tolerance scores for >14.7 million human amino acid sites. We calculated mean INDEL_i_ scores within the sliding windows used for DR score computation and compared these scores across regions with different DR values.

### Identifying sequence features associated with high INDEL mutation rates

Because of the high computational burden of related analyses in this section, we only focused on 1 bp and 2 bp INDEL subtypes. For each INDEL subtype, we considered the top 100,000 genomic positions with the highest MuRaL-indel-predicted mutation rates as hypermutated sites. We randomly sampled 100,000 positions as control sites. The method Integrated Gradients (IG)^31^ was applied to compute base-resolution contribution scores for each input sequence. The contribution of the central 200 bp region was quantified by comparing the gradient difference between the top 100,000 sites and the control group.

For motif discovery, we applied MoDISco^32^ to the ±100 bp flanking sequences of the top 100,000 sites, extracting and clustering positive seqlets into motifs. Motifs were trimmed by removing positions with information content < 0.1 from the position weight matrix (PWM). The contribution weight matrix (CWM) of each motif was obtained by multiplying the trimmed sequences with their corresponding contribution scores and averaging across seqlets. Motif-level contribution scores were then calculated by summing over all positions of the CWM. Candidate motifs were further filtered according to the following criteria:

1. the average observed rare variant density at genomic sites containing the motif within central 200bp was higher than the mean of the top 100,000 high mutation sites;
2. the number of motif occurrences in the analyzed regions exceeded 100;
3. the motif contribution score was higher than the median of all motifs.

Region enrichment analysis was performed using GAT^54^. Repeat annotations were obtained from RepeatMasker (https://hgdownload.soe.ucsc.edu/goldenPath/hg38/database/rmsk.txt.gz), Hip-STR^55^

(https://raw.githubusercontent.com/HipSTR-Tool/HipSTRreferences/master/human/GRCh38.hipstr_reference.bed.gz) and TRF^56^ (https://hgdownload.cse.ucsc.edu/goldenPath/hg38/bigZips/hg38.trf.bed.gz). The repeats from there sources were then merged. For the simple repeat category, elements with a repeat unit length of ≥6 (for 1 bp INDELs) and dinucleotide repeats with a unit length of ≥3 (for 2-bp INDELs) were further included as simple repeat elements in the annotation. To identify non-repeat motifs, we excluded all surveyed sites overlapping simple repeats within their ±5 bp flanking regions. Motif trimming and enrichment analyses were then repeated on the filtered set.

To assess whether motifs were the primary contributors to mutation, we compared the distribution of the motif occurrences, with the mean gradient profiles across the central 200 bp of the corresponding samples containing those seqlets. Gradient profiles were computed as the log-ratio of the central 200 bp gradients to those of randomly sampled control sites extended by 200 bp. Both motif occurrences and gradient profiles were normalized by Z-score.

### Additional analysis of the GTTACG motif

We first validated the hypermutability of the CTTACG motif using rare variant density and analyzed the average rare variant density of all 6-mers across the genome. The average rare variant density for a given *k*-mer was computed by the following formula:

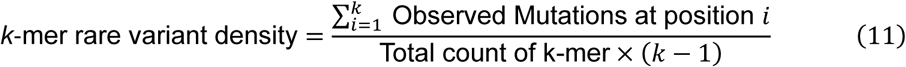

*k* is the length of the motif (e.g., *k* = 6 for a 6-mer). The position *i* represents the specific INDEL site within the motif. The index *i* = 1 denotes a mutation observed between the first and second base of the motif. *k-1* means *k-1* possible INDEL breakpoints in a *k*-mer.

To investigate the relationship between mutability and transcriptional context, we categorized CTTACG motifs based on their orientation relative to a gene’s transcribed strand. The average rare variant density was then calculated separately for motifs on the transcribed and non-transcribed strands. Next, we used the predicted average mutation rate (calculated using the Equation (11) as the observed rate) to partition all instances of the CTTACG motif into ten decile bins. Within each bin, we compared the average rare variant densities on the transcribed and non-transcribed strands. Moreover, we analyzed the enrichment of 5’ splice sites (5SS) within these decile bins. A 5SS was defined as the first base of an intron immediately downstream of an exon, based on the gene annotations. The proportion of motifs containing a 5SS was then calculated for each of the ten bins.

## Data availability

All the analyses in this study were based on published data. The predicted mutation rate maps for genomes of human, *M. mulatta*, *A. thaliana* and *D. melanogaster* are available at the ScienceDB repository: https://doi.org/10.57760/sciencedb.30830.

## Code availability

The MuRaL-indel model has been incorporated in the open-source MuRaL package, available at https://github.com/CaiLiLab/MuRaL.

## Acknowledgements

This work was supported by National Natural Science Foundation of China (32470690), State Key Laboratory of Biocontrol and Guangdong Provincial Key Laboratory for Aquatic Economic Animals.

## Author contributions

C.L. conceived and supervised the project. S.D. and H.S. developed the MuRaL-indel model and performed detailed analysis. All authors wrote the manuscript.

## References

1 Conrad, D. F. et al. Variation in genome-wide mutation rates within and between human families. Nat Genet 43, 712–714, doi:10.1038/ng.862 (2011).

2 Byrska-Bishop, M. et al. High-coverage whole-genome sequencing of the expanded 1000 Genomes Project cohort including 602 trios. Cell 185, 3426–3440 e3419, doi:10.1016/j.cell.2022.08.004 (2022).

3 Lin, M. et al. Effects of short indels on protein structure and function in human genomes. Sci Rep 7, 9313, doi:10.1038/s41598-017-09287-x (2017).

4 Riordan, J. R. et al. Identification of the cystic fibrosis gene: cloning and characterization of complementary DNA. Science 245, 1066–1073, doi:10.1126/science.2475911 (1989).

5 Gouw, S. C. et al. F8 gene mutation type and inhibitor development in patients with severe hemophilia A: systematic review and meta-analysis. Blood 119, 2922–2934, doi:10.1182/blood-2011-09-379453 (2012).

6 Wang, Y. & Obbard, D. J. Experimental estimates of germline mutation rate in eukaryotes: a phylogenetic meta-analysis. Evol Lett 7, 216–226, doi:10.1093/evlett/qrad027 (2023).

7 Kondrashov, A. S. & Rogozin, I. B. Context of deletions and insertions in human coding sequences. Hum Mutat 23, 177–185, doi:10.1002/humu.10312 (2004).

8 Kvikstad, E. M. & Duret, L. Strong heterogeneity in mutation rate causes misleading hallmarks of natural selection on indel mutations in the human genome. Mol Biol Evol 31, 23–36, doi:10.1093/molbev/mst185 (2014).

9 Mills, R. E. et al. An initial map of insertion and deletion (INDEL) variation in the human genome. Genome Res 16, 1182–1190, doi:10.1101/gr.4565806 (2006).

10 Kvikstad, E. M., Tyekucheva, S., Chiaromonte, F. & Makova, K. D. A macaque’s-eye view of human insertions and deletions: differences in mechanisms. PLoS Comput Biol 3, 1772–1782, doi:10.1371/journal.pcbi.0030176 (2007).

11 Maiolo, M. et al. ProPIP: a tool for progressive multiple sequence alignment with Poisson Indel Process. BMC Bioinformatics 22, 518, doi:10.1186/s12859-021-04442-8 (2021).

12 Iglhaut, C., Pecerska, J., Gil, M. & Anisimova, M. Please Mind the Gap: Indel-Aware Parsimony for Fast and Accurate Ancestral Sequence Reconstruction and Multiple Sequence Alignment Including Long Indels. Mol Biol Evol 41, doi:10.1093/molbev/msae109 (2024).

13 Clark, T. G. et al. Functional constraint and small insertions and deletions in the ENCODE regions of the human genome. Genome Biol 8, R180, doi:10.1186/gb-2007-8-9-r180 (2007).

14 Montgomery, S. B. et al. The origin, evolution, and functional impact of short insertion-deletion variants identified in 179 human genomes. Genome Res 23, 749–761, doi:10.1101/gr.148718.112 (2013).

15 Saitou, N. & Ueda, S. Evolutionary rates of insertion and deletion in noncoding nucleotide sequences of primates. Mol Biol Evol 11, 504–512, doi:10.1093/oxfordjournals.molbev.a040130 (1994).

16 Zhang, Z. & Gerstein, M. Patterns of nucleotide substitution, insertion and deletion in the human genome inferred from pseudogenes. Nucleic Acids Res 31, 5338–5348, doi:10.1093/nar/gkg745 (2003).

17 Fan, Y. et al. Patterns of insertion and deletion in Mammalian genomes. Curr Genomics 8, 370–378, doi:10.2174/138920207783406479 (2007).

18 Tanay, A. & Siggia, E. D. Sequence context affects the rate of short insertions and deletions in flies and primates. Genome Biol 9, R37, doi:10.1186/gb-2008-9-2-r37 (2008).

19 Carlson, J. et al. Extremely rare variants reveal patterns of germline mutation rate heterogeneity in humans. Nat Commun 9, 3753, doi:10.1038/s41467-018-05936-5 (2018).

20 Koh, G. C. C. et al. A redefined InDel taxonomy provides insights into mutational signatures. Nat Genet 57, 1132–1141, doi:10.1038/s41588-025-02152-y (2025).

21 Bethune, J., Kleppe, A. & Besenbacher, S. A method to build extended sequence context models of point mutations and indels. Nat Commun 13, 7884, doi:10.1038/s41467-022-35596-5 (2022).

22 Zhou, J. & Troyanskaya, O. G. Predicting effects of noncoding variants with deep learning-based sequence model. Nat Methods 12, 931–934, doi:10.1038/nmeth.3547 (2015).

23 Xu, S. et al. SSBlazer: a genome-wide nucleotide-resolution model for predicting single-strand break sites. Genome Biol 25, 46, doi:10.1186/s13059-024-03179-w (2024).

24 Benegas, G., Albors, C., Aw, A. J., Ye, C. & Song, Y. S. A DNA language model based on multispecies alignment predicts the effects of genome-wide variants. Nat Biotechnol, doi:10.1038/s41587-024-02511-w (2025).

25 Fang, Y., Deng, S. & Li, C. A generalizable deep learning framework for inferring fine-scale germline mutation rate maps. Nature Machine Intelligence 4, 1209–1223, doi:10.1038/s42256-022-00574-5 (2022).

26 Kull, M., Perello-Nieto, M., Kängsepp, M., Song, H. & Flach, P. Beyond temperature scaling: Obtaining well-calibrated multiclass probabilities with Dirichlet calibration. *arXiv preprint arXiv:1910.12656* (2019).

27 Chen, S. et al. A genomic mutational constraint map using variation in 76,156 human genomes. Nature 625, 92–100, doi:10.1038/s41586-023-06045-0 (2024).

28 Ding, W. et al. Adaptive functions of structural variants in human brain development. Sci Adv 10, eadl4600, doi:10.1126/sciadv.adl4600 (2024).

29 Halldorsson, B. V. et al. The sequences of 150,119 genomes in the UK Biobank. Nature 607, 732–740, doi:10.1038/s41586-022-04965-x (2022).

30 Topolska, M., Beltran, A. & Lehner, B. Deep indel mutagenesis reveals the impact of amino acid insertions and deletions on protein stability and function. Nat Commun 16, 2617, doi:10.1038/s41467-025-57510-5 (2025).

31. Sundararajan, M., Taly, A. & Yan, Q. Axiomatic Attribution for Deep Networks. doi:10.48550/arXiv.1703.01365 (2017).

32 Shrikumar, A. et al. Technical Note on Transcription Factor Motif Discovery from Importance Scores (TF-MoDISco) version 0.5.6.5. doi:10.48550/arXiv.1811.00416 (2020).

33 Chuzhanova, N. A., Anassis, E. J., Ball, E. V., Krawczak, M. & Cooper, D. N. Meta-analysis of indels causing human genetic disease: mechanisms of mutagenesis and the role of local DNA sequence complexity. Human Mutation 21, 28–44, doi:10.1002/humu.10146 (2003).

34 Cooper, D. N., Stenson, P. D. & Chuzhanova, N. A. The Human Gene Mutation Database (HGMD) and Its Exploitation in the Study of Mutational Mechanisms. Current Protocols in Bioinformatics 12, 1.13.11–11.13.20, doi:10.1002/0471250953.bi0113s12 (2005).

35 Kadri, N. K., Mapel, X. M. & Pausch, H. The intronic branch point sequence is under strong evolutionary constraint in the bovine and human genome. Communications Biology 4, 1206, doi:10.1038/s42003-021-02725-7 (2021).

36 He, K., Zhang, X., Ren, S. & Sun, J. in 2016 *IEEE Conference on Computer Vision and Pattern Recognition (CVPR)*. 770–778.

37 Paszke, A. et al. Pytorch: An imperative style, high-performance deep learning library. Advances in neural information processing systems 32, 8026–8037 (2019).

38. Dale, R. K., Pedersen, B. S. & Quinlan, A. R. Pybedtools: a flexible Python library for manipulating genomic datasets and annotations. Bioinformatics 27, 3423–3424, doi:10.1093/bioinformatics/btr539 (2011).

39. Kingma, D. P. & Ba, J. Adam: A method for stochastic optimization. arXiv preprint arXiv:1412.6980 (2014).

40. Liaw, R., et al. Tune: A research platform for distributed model selection and training. arXiv preprint arXiv:1807.05118 (2018).

41 Zhu, Y. O., Sherlock, G. & Petrov, D. A. Extremely Rare Polymorphisms in Saccharomyces cerevisiae Allow Inference of the Mutational Spectrum. PLoS Genet 13, e1006455, doi:10.1371/journal.pgen.1006455 (2017).

42 Agarwal, I. & Przeworski, M. Signatures of replication timing, recombination, and sex in the spectrum of rare variants on the human X chromosome and autosomes. Proc Natl Acad Sci U S A 116, 17916–17924, doi:10.1073/pnas.1900714116 (2019).

43 Zhao, G. et al. Gene4Denovo: an integrated database and analytic platform for de novo mutations in humans. Nucleic Acids Res 48, D913–D926, doi:10.1093/nar/gkz923 (2020).

44 Jonsson, H. et al. Parental influence on human germline de novo mutations in 1,548 trios from Iceland. Nature 549, 519–522, doi:10.1038/nature24018 (2017).

45 Yuen, R. et al. Whole genome sequencing resource identifies 18 new candidate genes for autism spectrum disorder. Nat Neurosci 20, 602–611, doi:10.1038/nn.4524 (2017).

46 An, J. Y. et al. Genome-wide de novo risk score implicates promoter variation in autism spectrum disorder. Science 362, doi:10.1126/science.aat6576 (2018).

47 Besenbacher, S. et al. Novel variation and de novo mutation rates in population-wide de novo assembled Danish trios. Nat Commun 6, 5969, doi:10.1038/ncomms6969 (2015).

48 Vasimuddin, M., Misra, S., Li, H. & Aluru, S. in 2019 IEEE International Parallel and Distributed Processing Symposium (IPDPS). 314–324.

49 Consortium, T. G. 1,135 Genomes Reveal the Global Pattern of Polymorphism in Arabidopsis thaliana. Cell 166, 481–491, doi:10.1016/j.cell.2016.05.063 (2016).

50 Huang, W. et al. Natural variation in genome architecture among 205 Drosophila melanogaster Genetic Reference Panel lines. Genome Res 24, 1193–1208, doi:10.1101/gr.171546.113 (2014).

51 Frankish, A. et al. GENCODE: reference annotation for the human and mouse genomes in 2023. Nucleic Acids Res 51, D942–D949, doi:10.1093/nar/gkac1071 (2023).

52 Ramirez, F. et al. deepTools2: a next generation web server for deep-sequencing data analysis. Nucleic Acids Res 44, W160–165, doi:10.1093/nar/gkw257 (2016).

53 Berrio, A., Haygood, R. & Wray, G. A. Identifying branch-specific positive selection throughout the regulatory genome using an appropriate proxy neutral. BMC Genomics 21, 359, doi:10.1186/s12864-020-6752-4 (2020).

54 Heger, A., Webber, C., Goodson, M., Ponting, C. P. & Lunter, G. GAT: a simulation framework for testing the association of genomic intervals. Bioinformatics 29, 2046–2048, doi:10.1093/bioinformatics/btt343 (2013).

55 Willems, T. et al. Genome-wide profiling of heritable and de novo STR variations. Nature Methods 14, 590–592, doi:10.1038/nmeth.4267 (2017).

56 Benson, G. Tandem repeats finder: a program to analyze DNA sequences. Nucleic Acids Research 27, 573–580, doi:10.1093/nar/27.2.573 (1999).

