## Supplementary Figures and Tables for "A deep learning framework for building INDEL mutation rate maps"

**This document contains Supplementary Figures 1-26 and Supplementary Tables 1-10.**

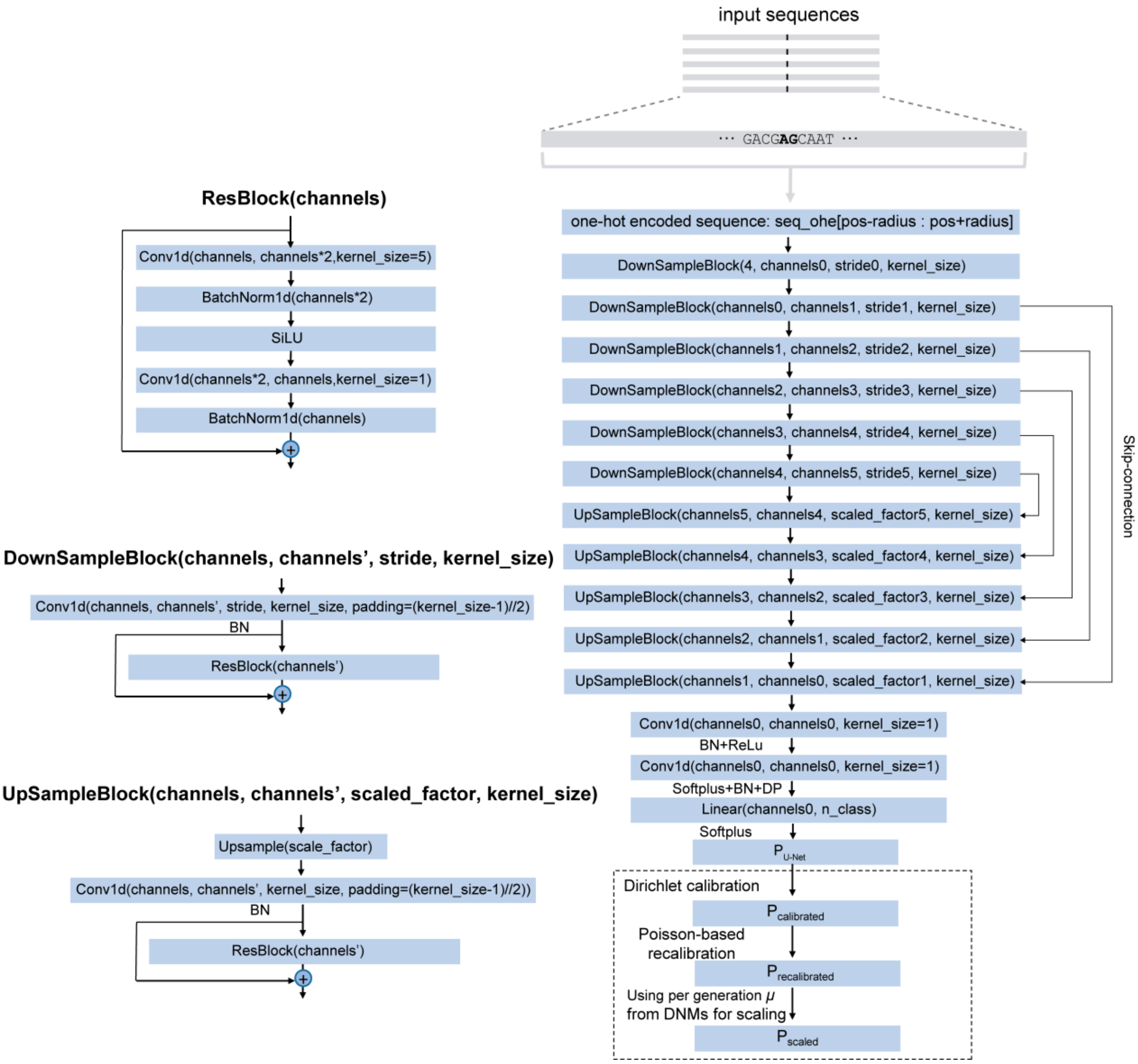


**Supplementary Fig. 1 Illustration for MuRaL-indel layers.** The figure depicts the encoder-decoder network architecture in detail. The encoder (downsampling path) comprises 6 DownSampleBlocks that progressively extract multi-scale features through strided convolution and residual blocks. The decoder contains 5 UpSampleBlocks that restore spatial resolution via upsampling and convolution operations. Features from corresponding scales are fused through skip connections. Each ResBlock consists of two convolutional layers with batch normalization and SiLU activation. Network outputs are processed through convolutional layers, fully connected layer, and Softplus activation, followed by Dirichlet calibration to produce final calibrated mutation rate predictions.

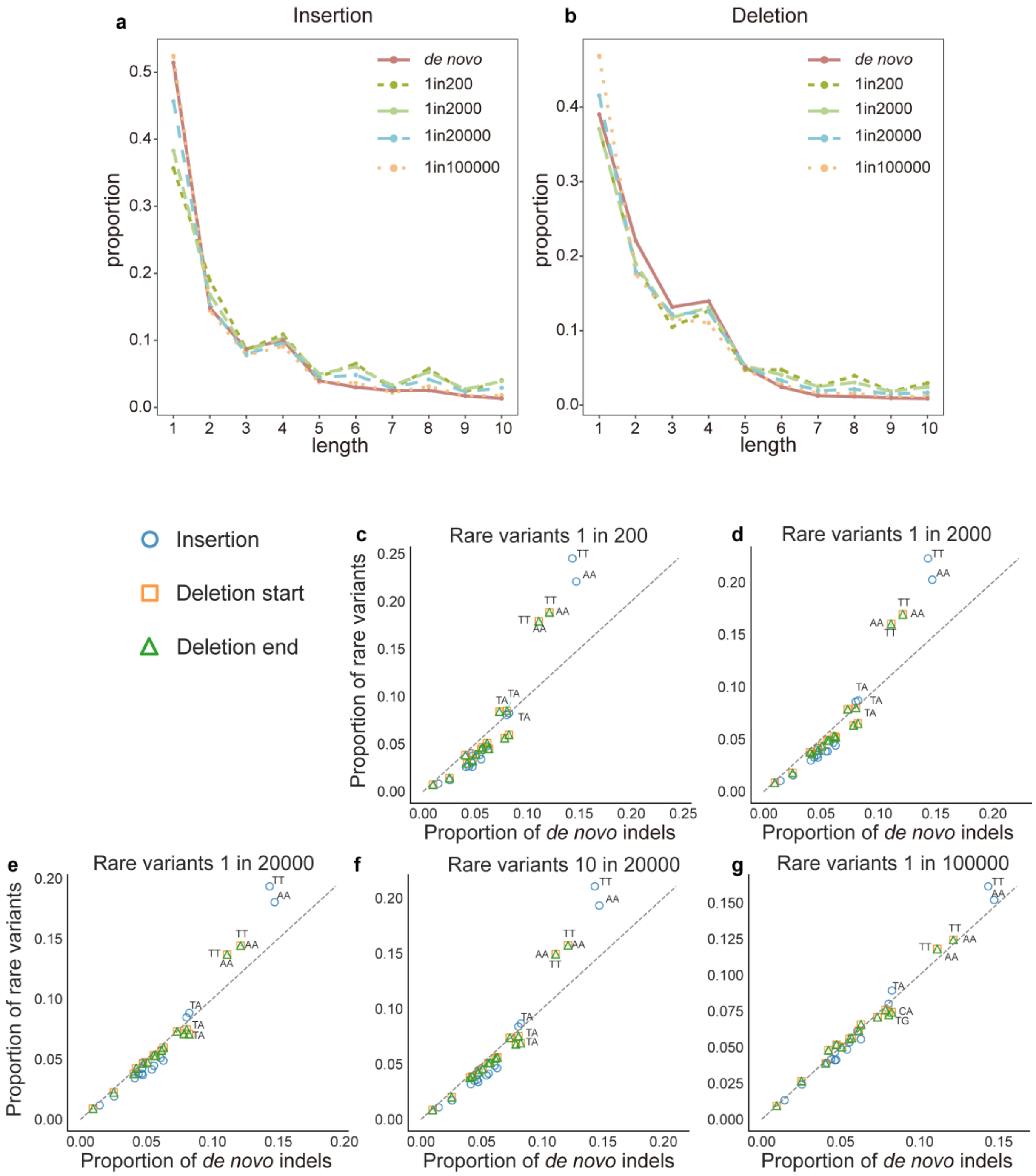


**Supplementary Fig. 2 Comparison of 2-mer mutation spectra of rare variants from different downsampled datasets with *de novo* INDELs.** (**a-b**) Distribution of *de novo* indels and various low-frequency INDELs across 1-10 bp lengths. The **a** shows insertions, and the **b** shows deletions. (**c-g**) Distribution of 1-bp INDELs in 2-mers for 1in200, 1in2000, 1in20000, 10in20000 and 1in100000 INDELs and *de novo* INDELs. The x-axis shows the proportion of different 2-mers in *de novo* INDELs, and the y-axis shows the proportion of different 2-mers in other low-frequency INDELs. The dashed lines represent the linear fittings of other 2-mer types that do not contain A and T.

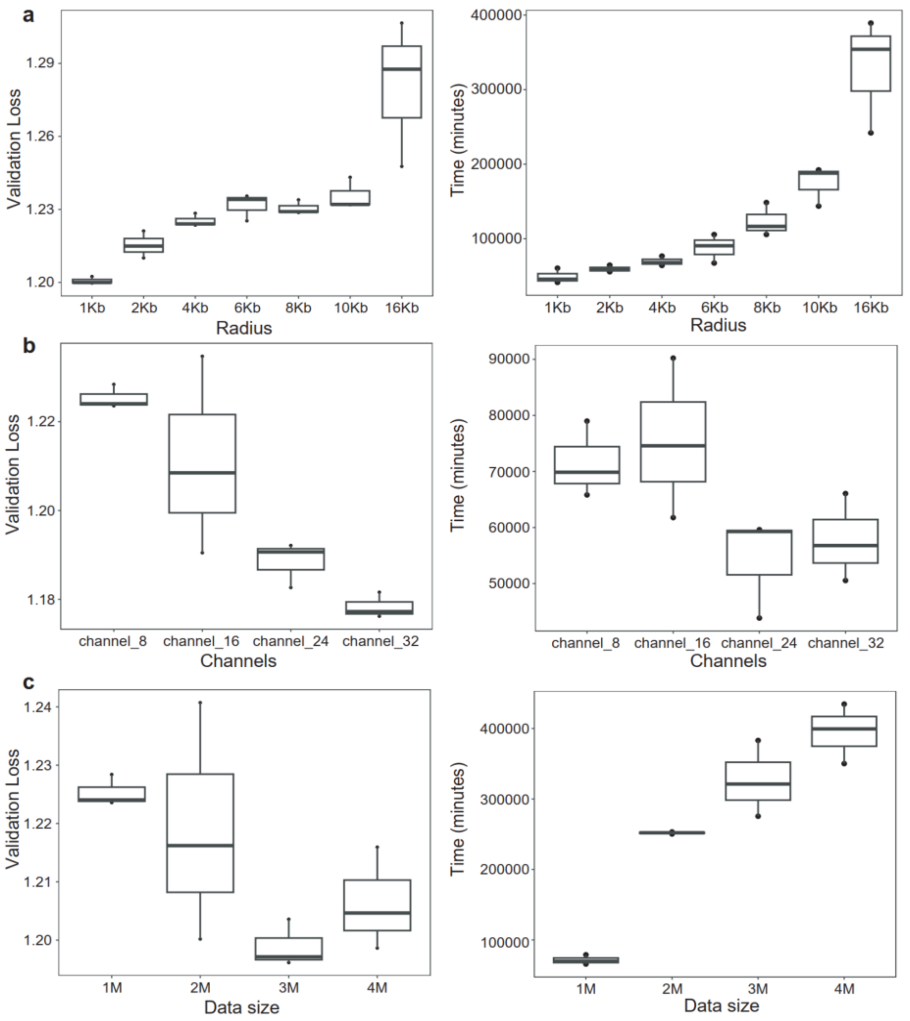


**Supplementary Fig. 3 Evaluation of models with different model configurations.** Box plots comparing validation loss (left panels) and training time cost (right panels) across three key hyperparameters. Each row represents a different optimization parameter: sequence radius, first-CNN-layer channel size, and training mutation dataset size (top to bottom). Left panels show the minimum validation loss achieved across three independent trials; right panels display the distribution of training time for each trial. (**a**) Validation loss increases with radius size, whereas larger radius values demonstrate improved *k*-mer and regional correlation patterns. Balancing prediction accuracy and computational efficiency, the radius of 4Kb was selected. (**b**) Validation loss decreases monotonically with increasing channel numbers, total training time does not increase proportionally owing to earlier convergence with higher model capacity. However, channel expansion substantially increases GPU memory and the model size. Considering the trade-off between model performance and resource constraints, channel_8 was adopted for the final model. (**c**) Training data sizes with 3M and 4M training mutations achieve markedly lower validation losses compared with smaller configurations. Combined with *k*-mer and regional correlation analysis, a size of 4M was selected.

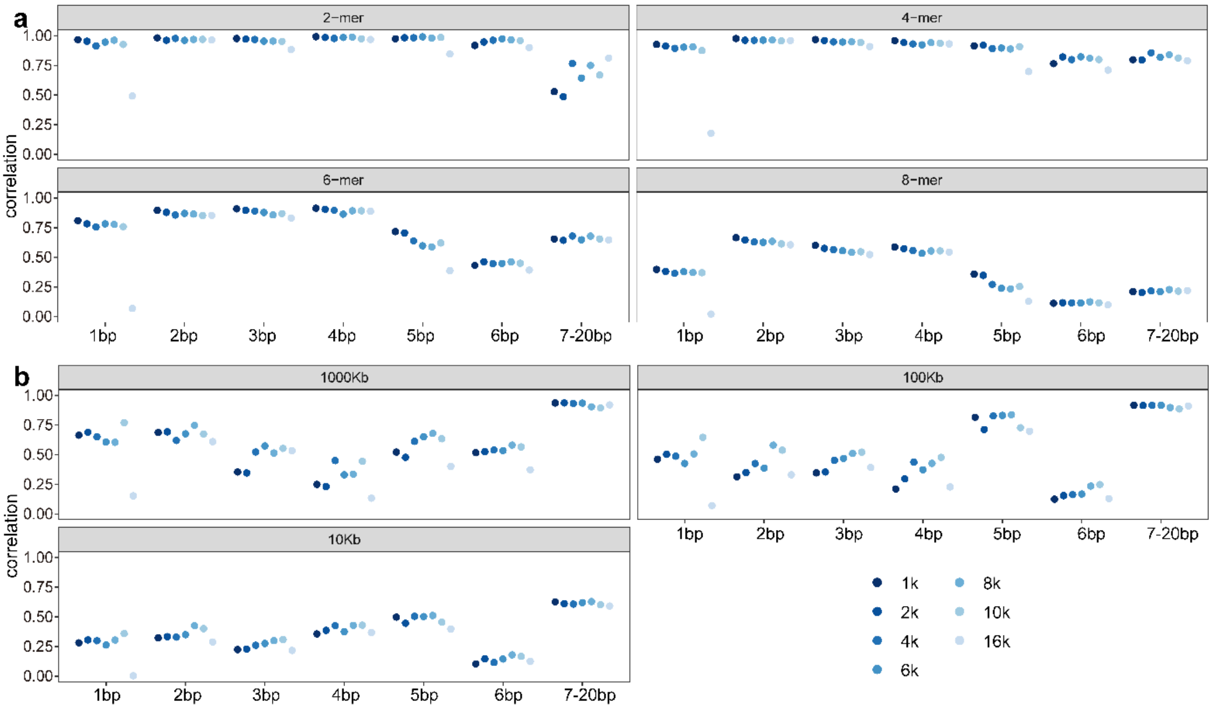


**Supplementary Fig. 4 Effect of sequence radius on the performance of the deletion start model.** (**a**) Effect of sequence radius on correlations between predicted and observed 2-, 4-, 6-, and 8-mer mutation rates, stratified by INDEL length. (**b**) Effect of sequence radius on correlations between predicted and observed regional mutation rates across genomic bins of 1Mb, 100Kb, and 10Kb, stratified by INDEL length. All correlations were based on Pearson's correlation tests.

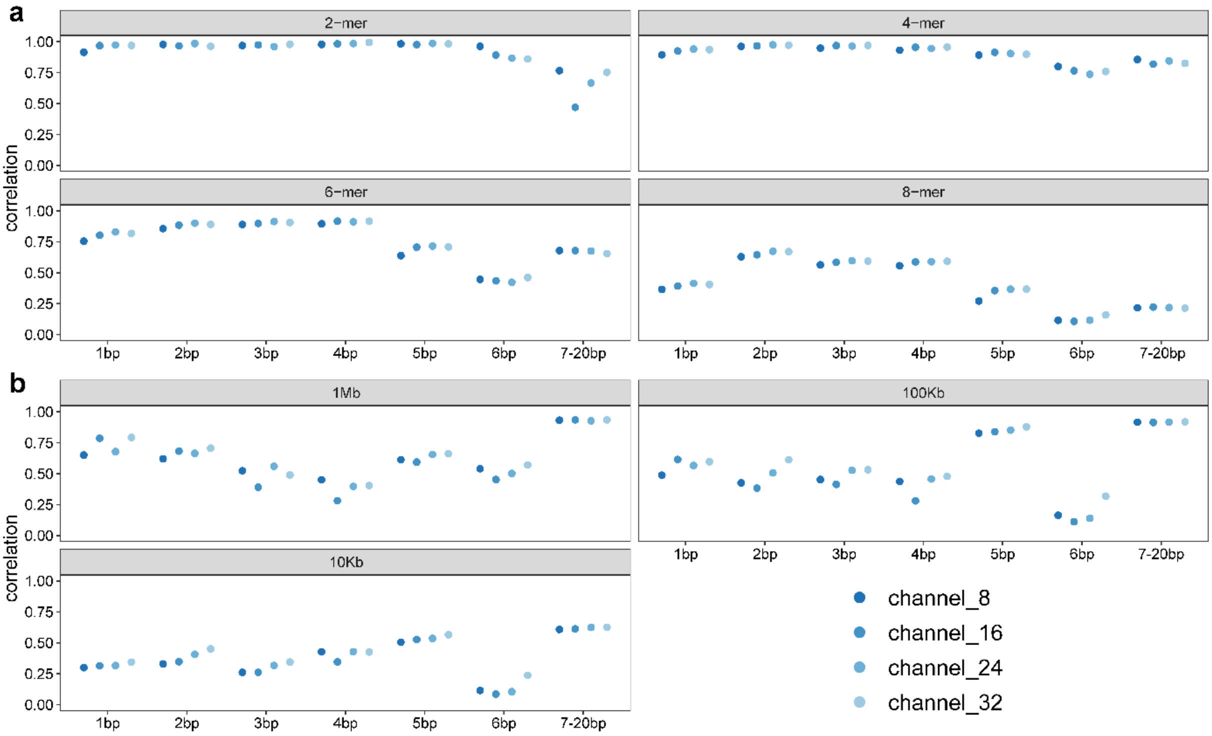


**Supplementary Fig. 5 Effect of channel size on the performance of the deletion start model. (a**) Effect of channel size on correlations between predicted and observed 2-, 4-, 6-, and 8-mer mutation rates, stratified by INDEL length. (**b**) Effect of channel size on correlations between predicted and observed regional mutation rates across genomic bins of 1Mb, 100Kb, and 10Kb, stratified by INDEL length. All correlations were based on Pearson's correlation tests.

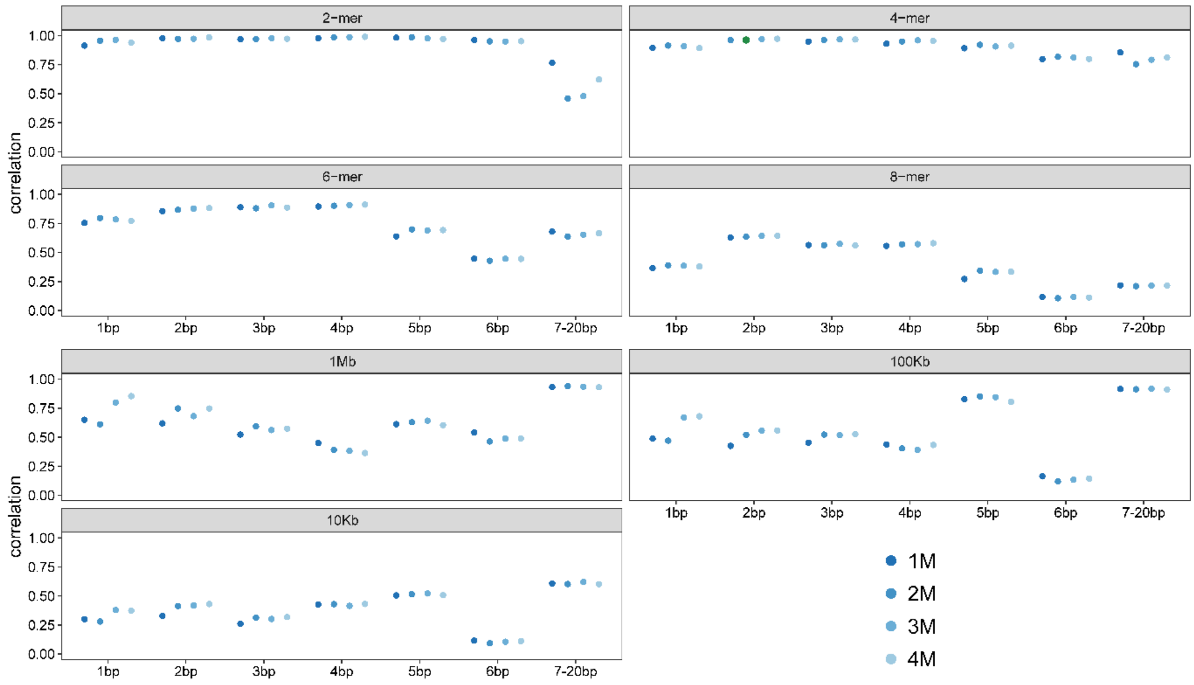


**Supplementary Fig. 6 Effect of training mutation data size on the performance of the deletion start model. (a**) Effect of data size on correlations between predicted and observed 2-, 4-, 6-, and 8-mer mutation rates, stratified by INDEL length. (**b**) Effect of data size on correlations between predicted and observed regional mutation rates across genomic bins of 1Mb, 100Kb, and 10Kb, stratified by INDEL length. All correlations were based on Pearson's correlation tests.

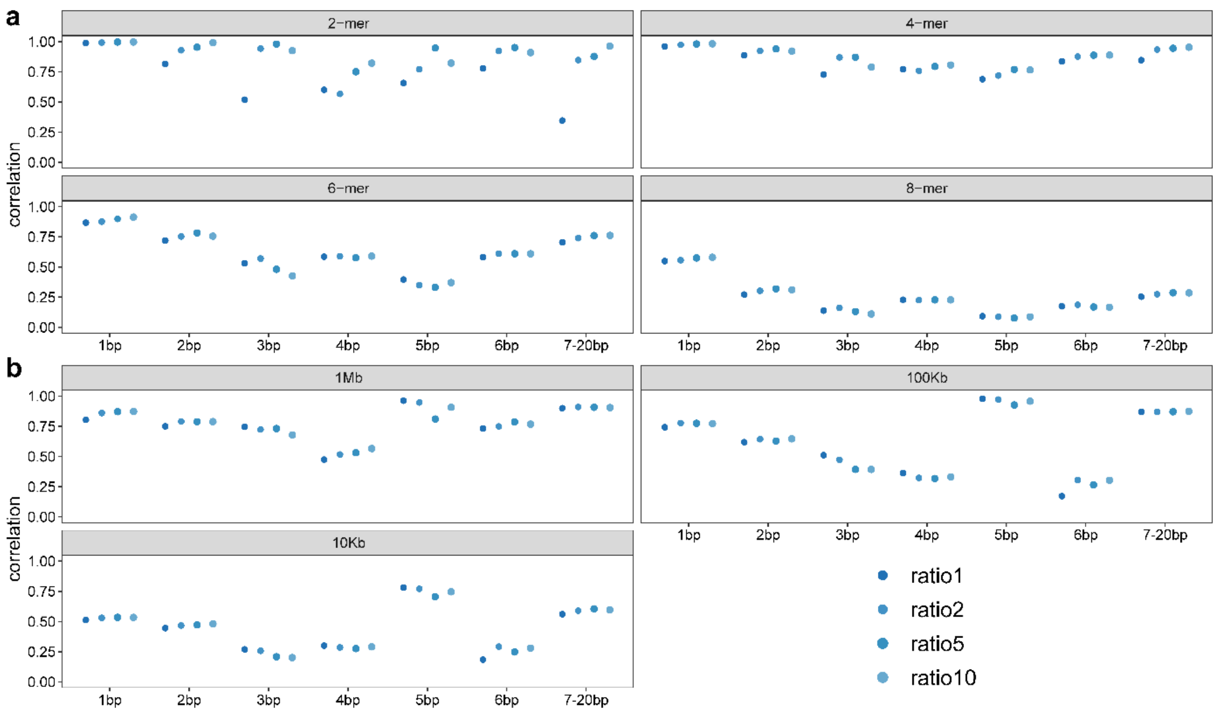


**Supplementary Fig. 7 Effect of training sample ratio (non-mutated to mutated) on the performance of the insertion model. (a**) Effect of training sample ratio on correlations between predicted and observed 2-, 4-, 6-, and 8-mer mutation rates, stratified by INDEL length. We fixed the number of mutated sites (1 million) and used different numbers of non-mutated sites. (**b**) Effect of sample ratio on correlations between predicted and observed regional mutation rates across genomic bins of 1Mb, 100Kb, and 10Kb, stratified by INDEL length. All correlations were based on Pearson's correlation tests. An optimal configuration was observed at a sample ratio of 1:10, which enhanced overall model performance.

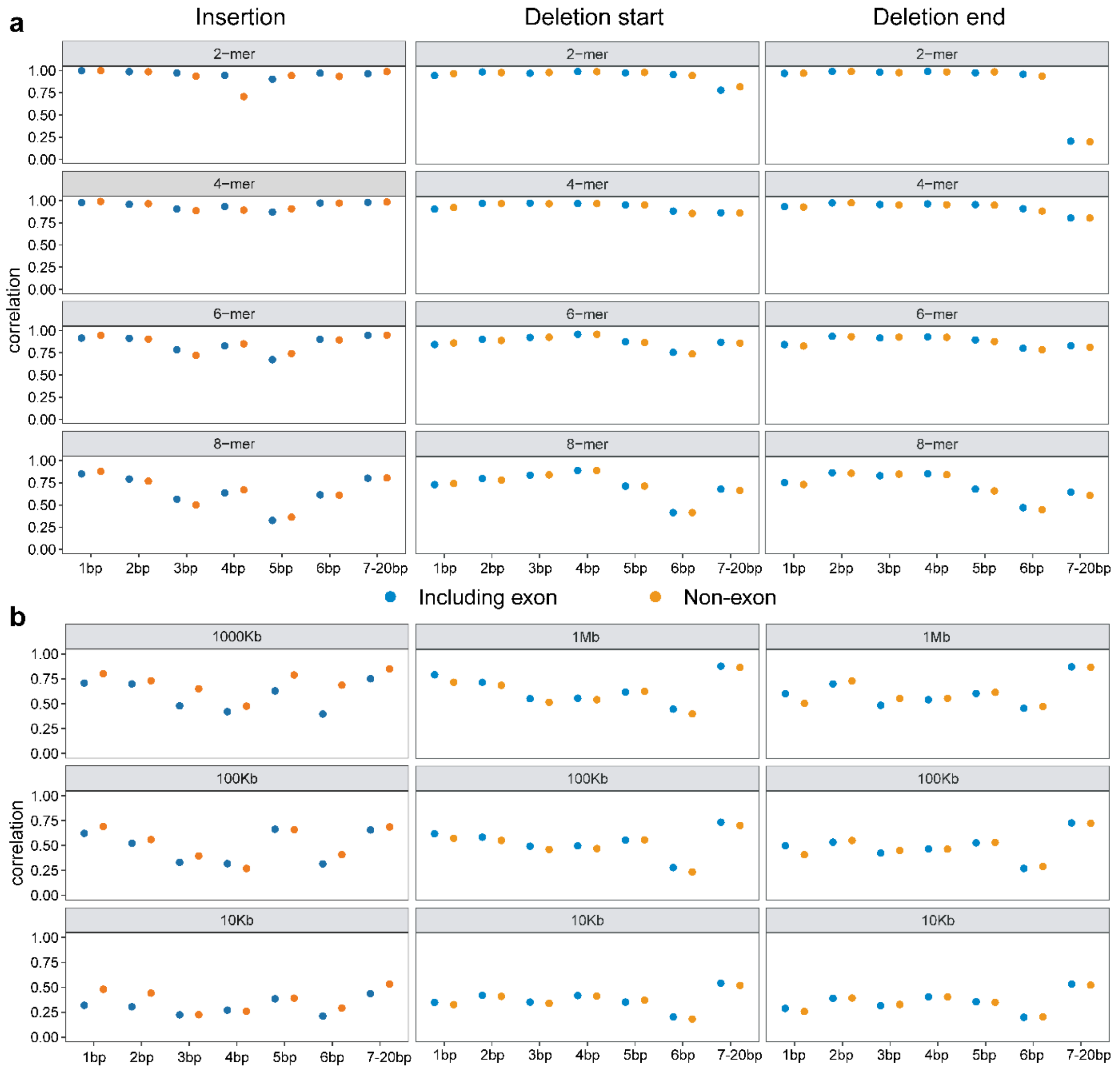


**Supplementary Fig. 8 Effect of exonic mutations on the performance of models. (a)** Effect of exonic mutations on correlations between predicted and observed 2-, 4-, 6-, and 8-mer mutation rates, stratified by INDEL length. (**b**) Effect of exonic mutation on correlations between predicted and observed regional mutation rates across genomic bins of 1Mb, 100Kb, and 10Kb, stratified by INDEL length. All correlations were based on Pearson's correlation tests. Overall, the exon-containing model and the exon-excluding model have similar *k*-mer correlation and regional correlation values. Considering that generally INDELs are subject to greater selection pressure, we used the human genome data without exons for training.

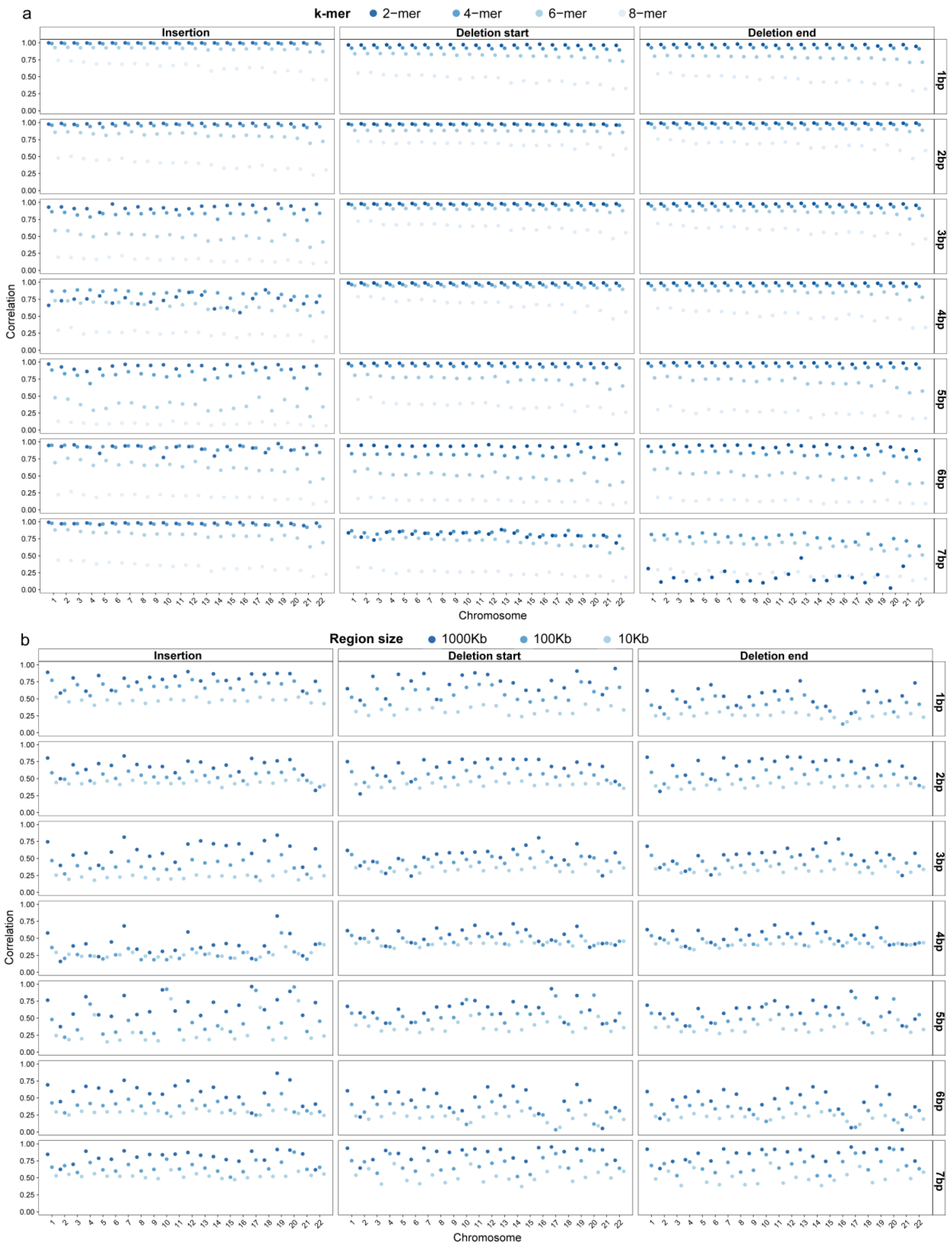


**Supplementary Fig. 9 *K*-mer and regional mutation rate correlations of the three MuRaL-indel models across autosomes.** (**a**) 2-, 4-, 6-, and 8-mer mutation rate correlations across autosomes for three INDEL models, stratified by INDEL length. (**b**) Regional mutation rate correlations across genomic bins of 1Mb, 100Kb, and 10Kb for three INDEL models, stratified by mutation type in each autosome. All correlations were based on Pearson's correlation tests.

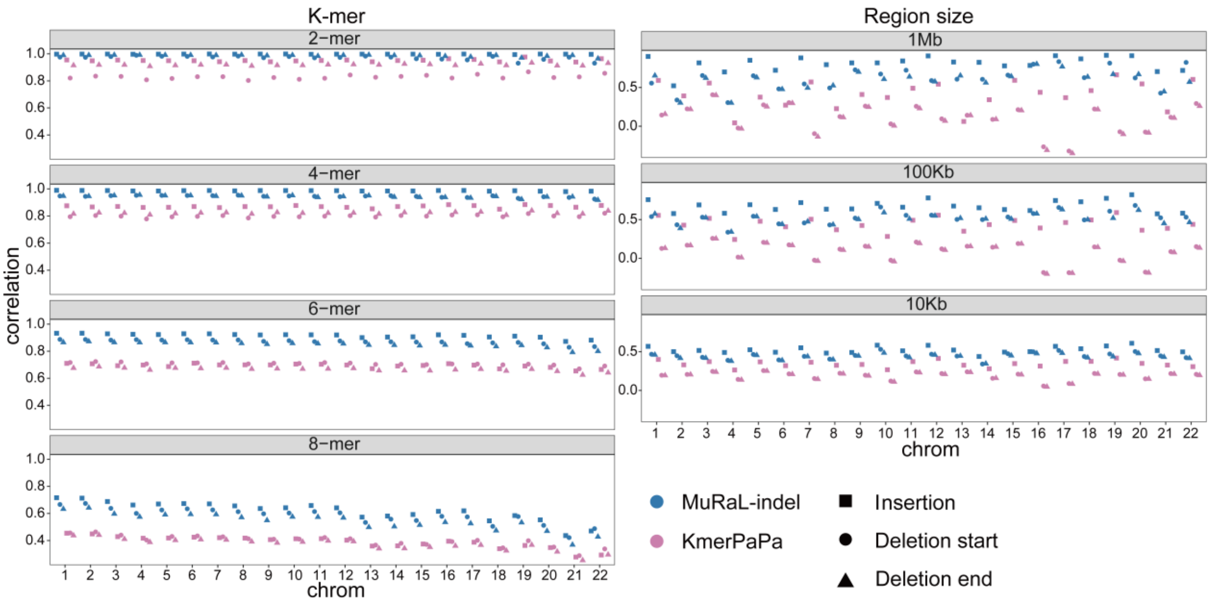


**Supplementary Fig. 10 Comparison of *k*-mer and regional mutation rate correlations between MuRaL-indel and kmerPaPa for three INDEL models across autosomes.** All correlations were based on Pearson's correlation tests.

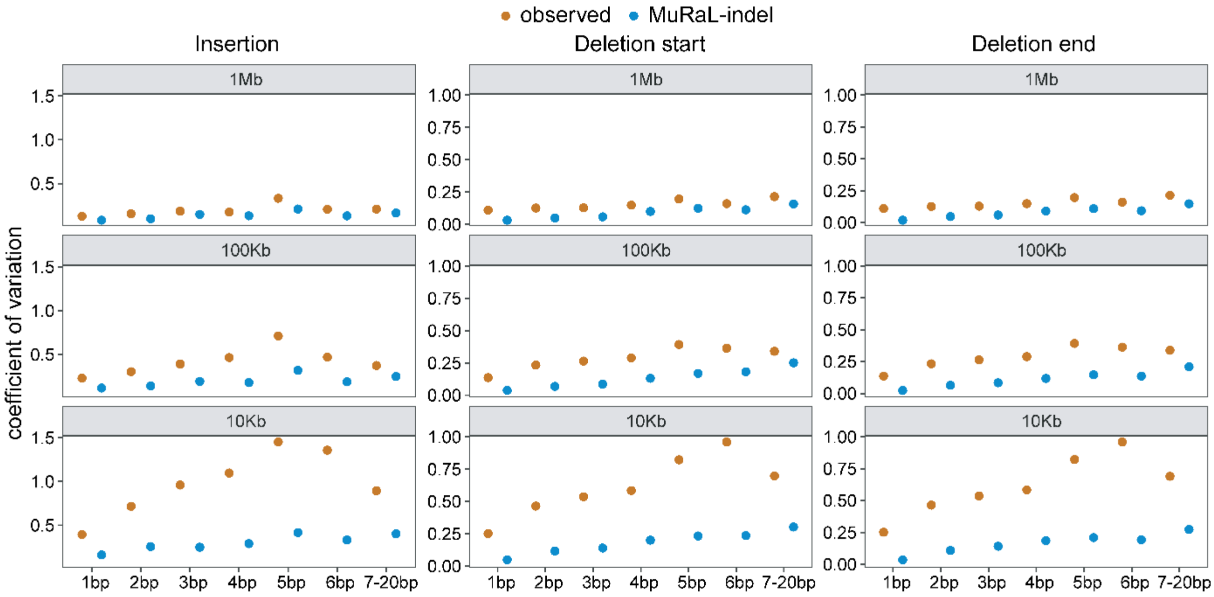


**Supplementary Fig. 11 Comparison of coefficient of variation (CV) of observed and predicted mutation rates for three INDEL models.** Brown symbols represent CV values derived from observed mutations (1in100000 rare variants), and blue symbols represent CV values from MuRaL-indel predicted mutation rates.

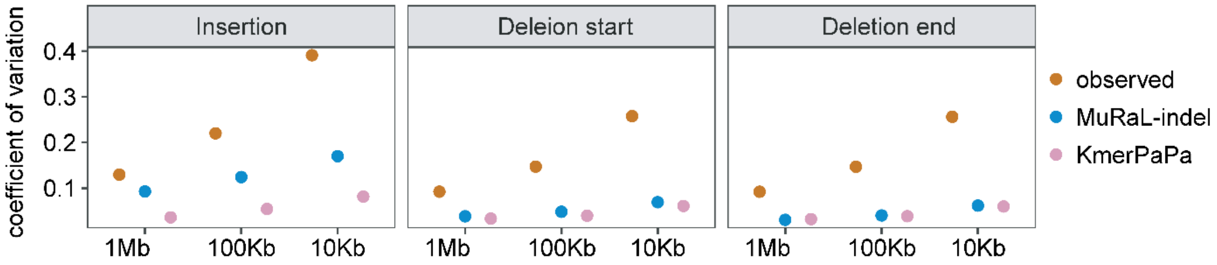


**Supplementary Fig. 12 Comparison of the coefficient of variation (CV) in regional mutation rates among observed, MuRaL-indel and kmerPaPa datasets for three INDEL models.** Average mutation rates were computed with bin sizes of 1Mb, 100Kb and 10Kb, and the CV was calculated separately at each scale. Brown symbols represent CV values derived from observed mutations (1in100000 subsample), blue symbols represent CV values from MuRaL-indel predicted mutation rates, and red symbols represent those from kmerPaPa-predicted mutation rates.

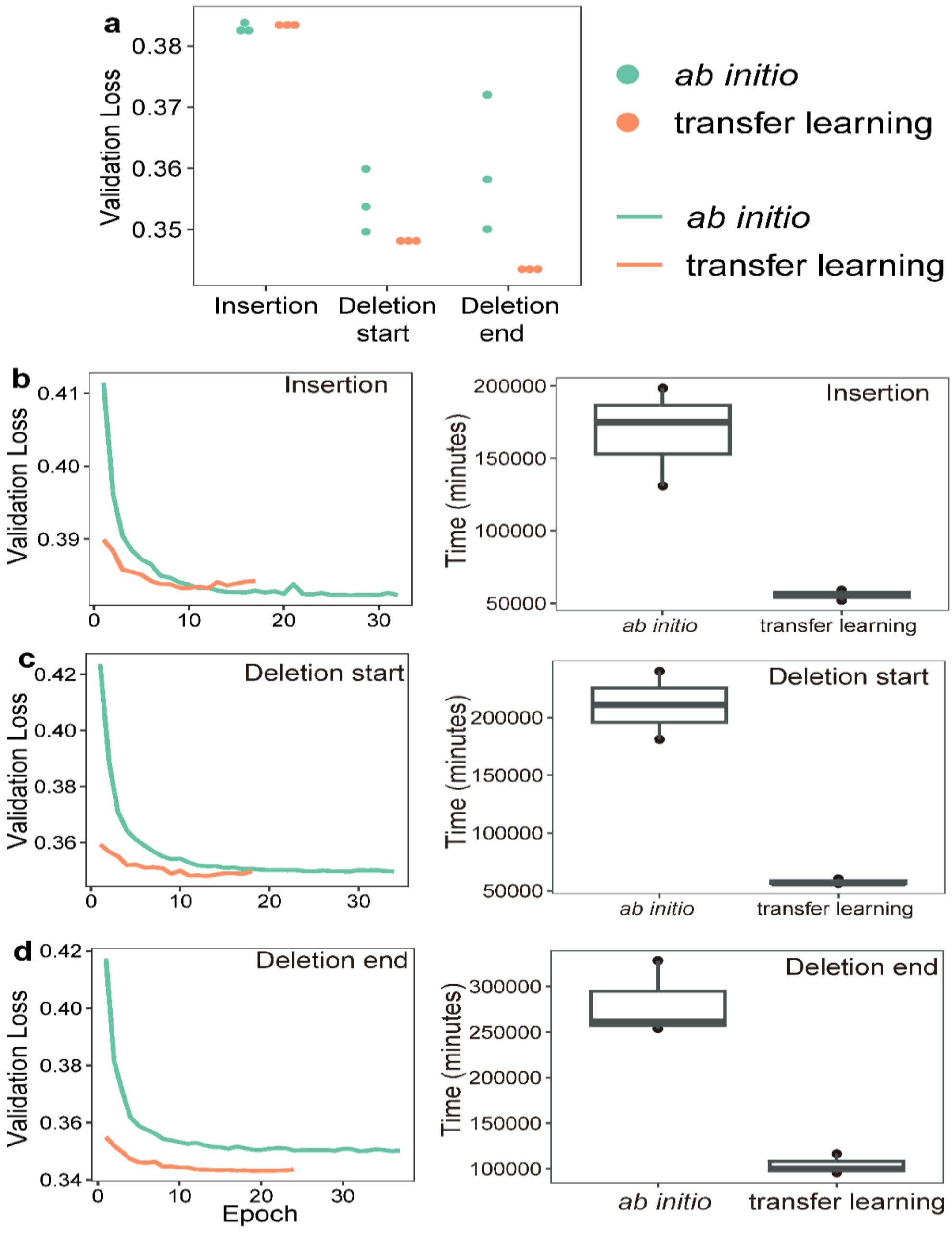


**Supplementary Fig. 13 Comparison of validation loss and training time between *ab initio* and transfer learning models for *M. mulatta*.** (**a**) Minimum validation loss achieved by *ab initio* and transfer learning models across autosomes in the *M. mulatta* genome. Each model was trained in three independent trials. (**b-d**) For each mutation type shown in **a**, the left panels depict the trajectories of validation loss across training epochs for the models achieving the minimum validation loss, with green representing *ab initio* models and orange representing transfer learning models. The right panels show box plots of training time per independent trial (in minutes). In the box plots, the center line indicates the median, boxes represent the interquartile range (IQR), and whiskers extend to the most extreme data points within 1.5× IQR.

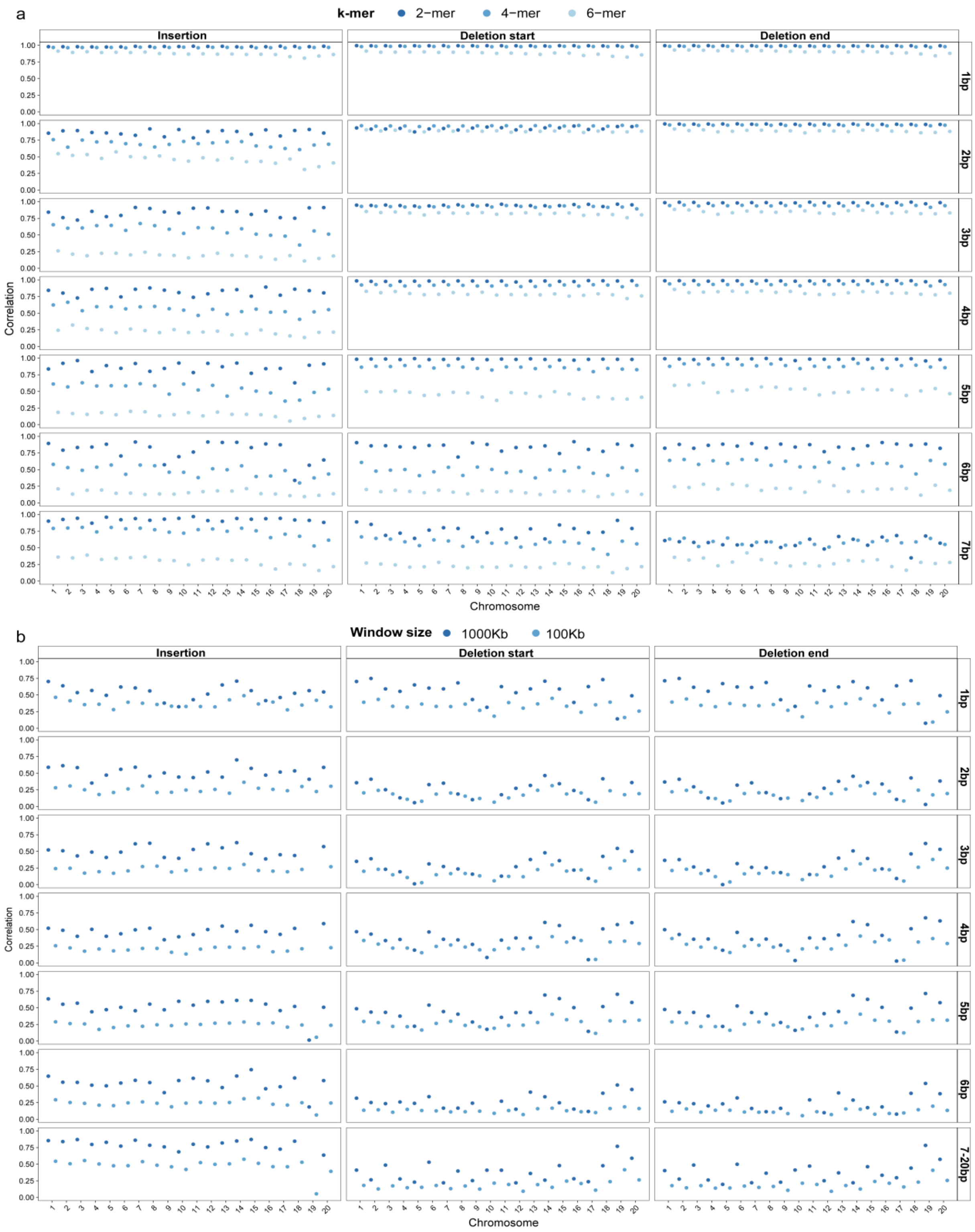


**Supplementary Fig. 14 *K*-mer and regional mutation rate correlations for three transfer learning MuRaL-indel models across autosomes in *M. mulatta*. (a**) 2-, 4-, 6-, and 8-mer mutation rate correlations across autosomes for three INDEL models, stratified by INDEL length. (**b)** Regional mutation rate correlations across genomic bins of 1Mb, 100Kb, and 10Kb for three INDEL models, stratified by mutation type in each autosome. All correlations were based on Pearson's correlation tests.

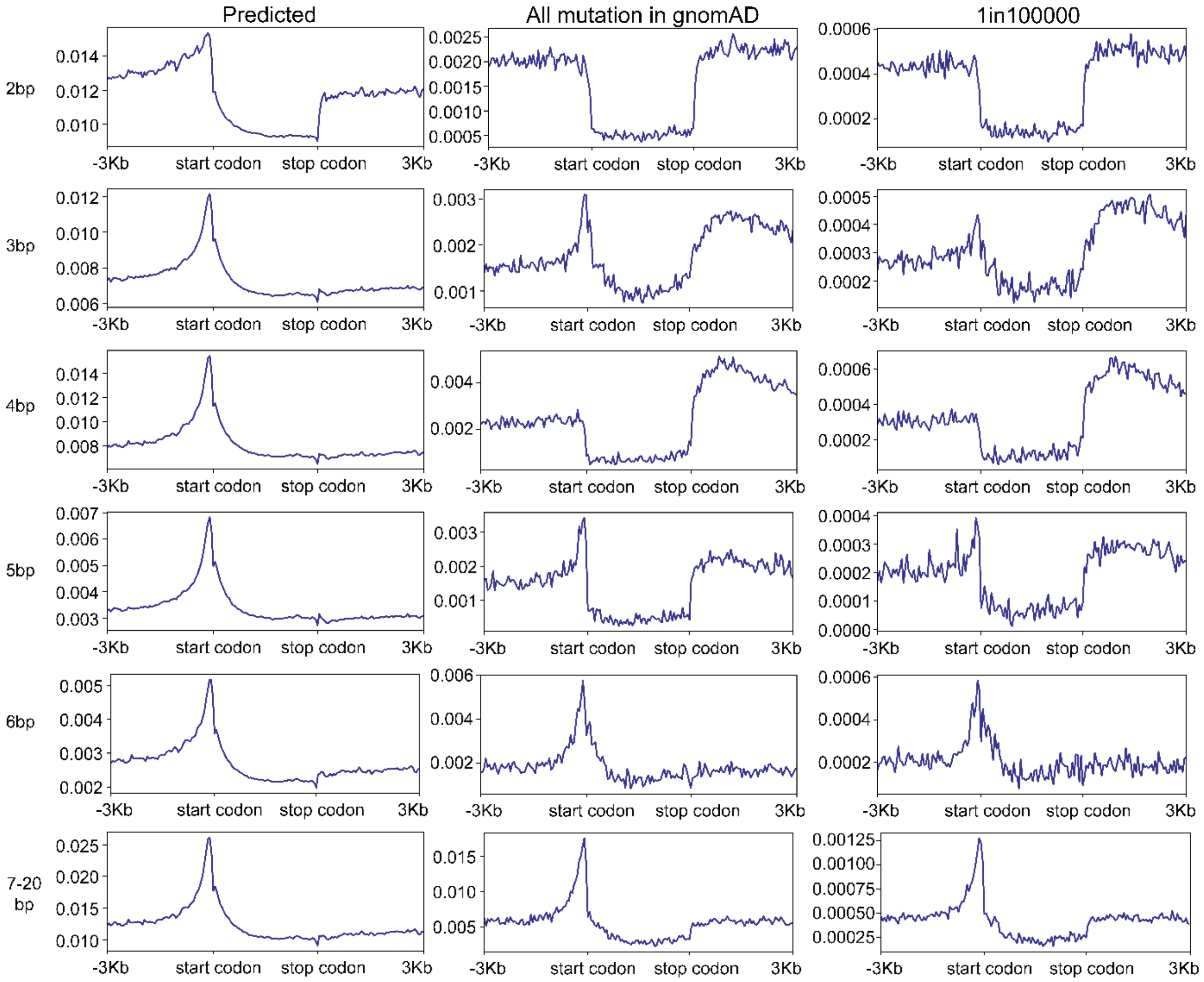


**Supplementary Fig. 15 The mutation rate patterns of insertions in human coding genes and their upstream and downstream regions.** This figure shows patterns for insertions of 2~20 bp, as the data of 1-bp insertions has been shown in the Fig. 4. The left panels depict the mutation rate predictions (non-scaled) for different insertion types in human coding genes and their upstream and downstream regions, as predicted by the MuRaL-indel model; The middle panels depict the variant distribution of different insertion types in human coding genes and their upstream and downstream regions, as recorded in the gnomAD database; The right panels depict the variant distribution of different insertion types with a very low frequency (1in100000) in human coding genes and their upstream and downstream regions, as recorded in the gnomAD database.

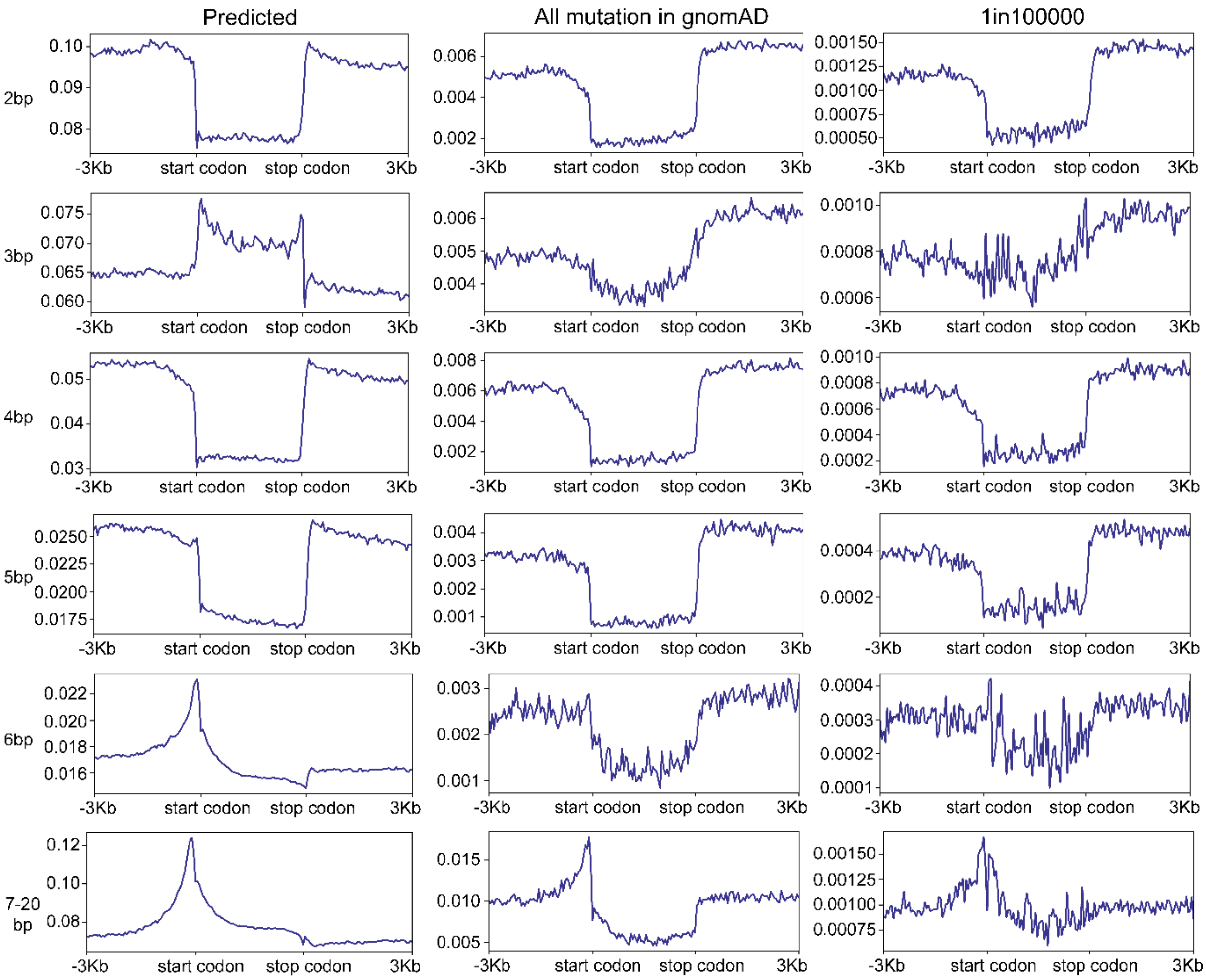


**Supplementary Fig. 16 The mutation rate patterns of deletion starts in human coding genes and their upstream and downstream regions.** This figure shows patterns for deletions of 2~20 bp, as the data of 1-bp insertions have been shown in the Fig. 4. The left panels depict the mutation rate predictions (non-scaled) for different deletion types in human coding genes and their upstream and downstream regions, as predicted by the MuRaL-indel model; The middle panels depict the variant distribution of different deletion types in human coding genes and their upstream and downstream regions, as recorded in the gnomAD database; The right panels depict the variant distribution of different deletion types with a very low frequency (1in100000) in human coding genes and their upstream and downstream regions, as recorded in the gnomAD database.

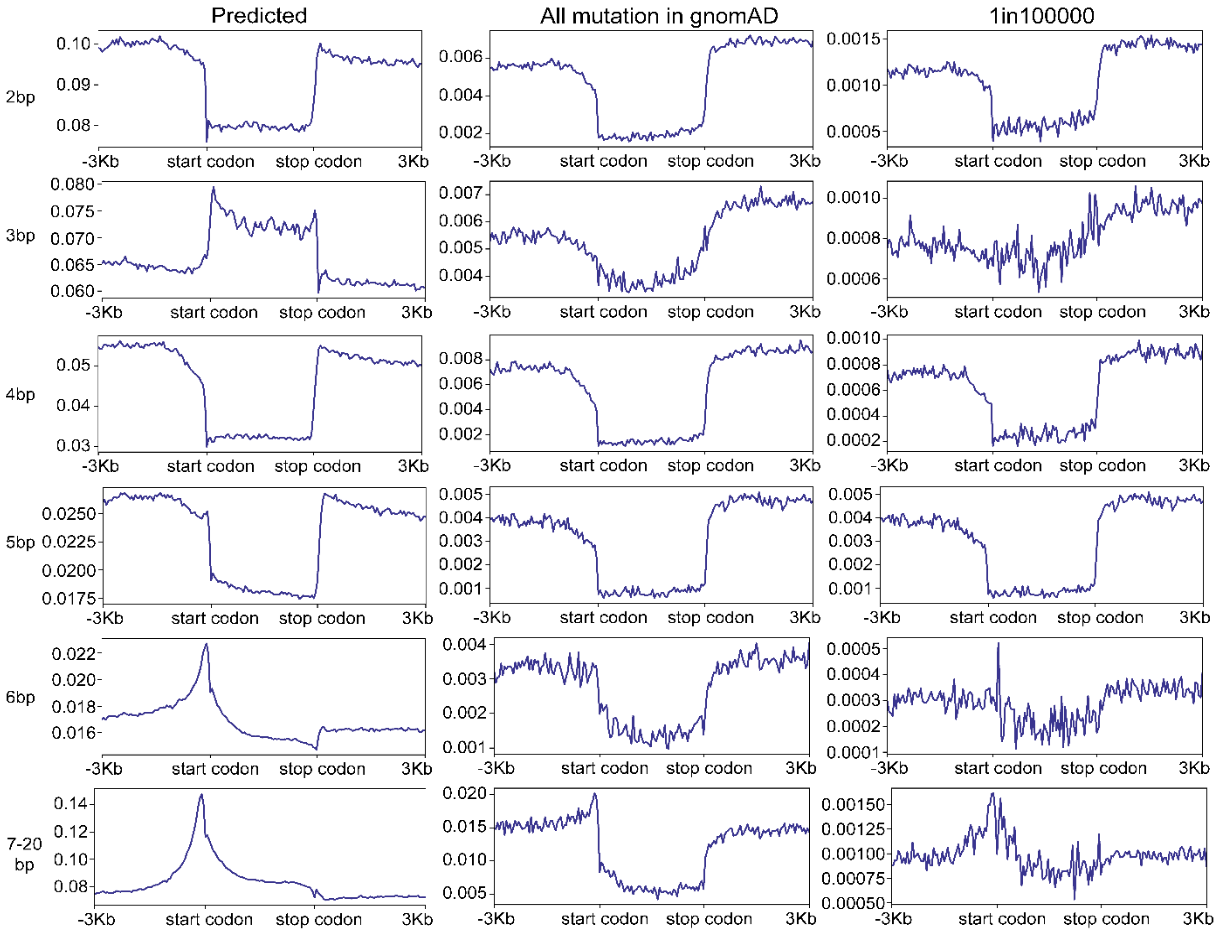


**Supplementary Fig. 17 The mutation rate patterns of deletion ends in human coding genes and their upstream and downstream regions.** This figure shows patterns for deletions of 2~20 bp, as the data of 1-bp insertions has been shown in the Fig. 4. The left panels depict the mutation rate predictions (non-scaled) for different deletion types in human coding genes and their upstream and downstream regions, as predicted by the MuRaL-indel model; The middle panels depict the variant distribution of different deletion types in human coding genes and their upstream and downstream regions, as recorded in the gnomAD database; The right panels depict the variant distribution of different deletion types with a very low frequency (1in100000) in human coding genes and their upstream and downstream regions, as recorded in the gnomAD database.

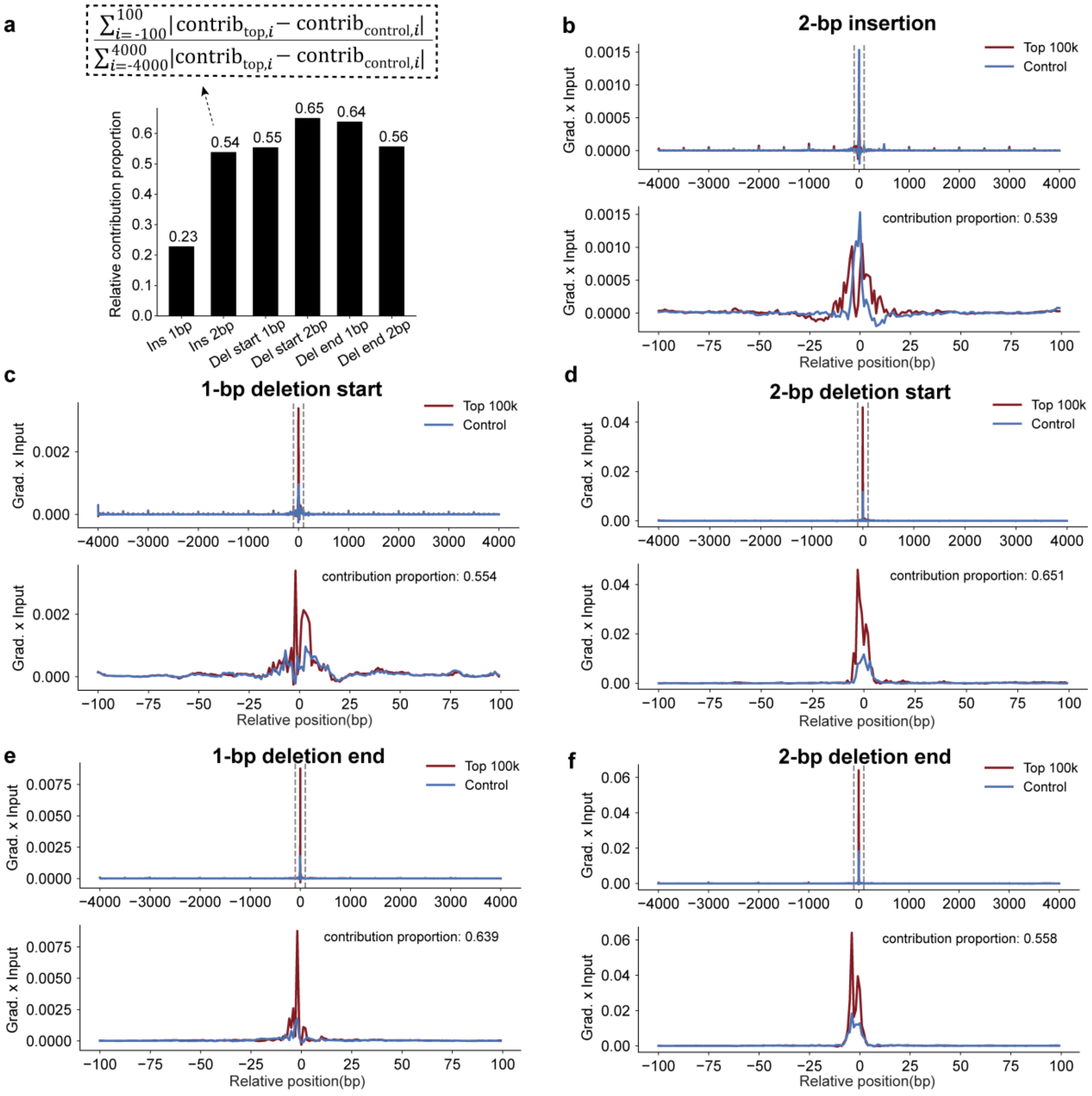


**Supplementary Fig. 18 Base-resolution contribution profiles for 1-bp and 2-bp INDELs.** (**a**) The formula (shown in the dashed box) defines the relative contribution proportion from the central 200bp region within each 8kb context, "contrib" denotes the product of the gradient and the input (gradient × input). (**b–f**) Average contribution of each base within the central 200bp.

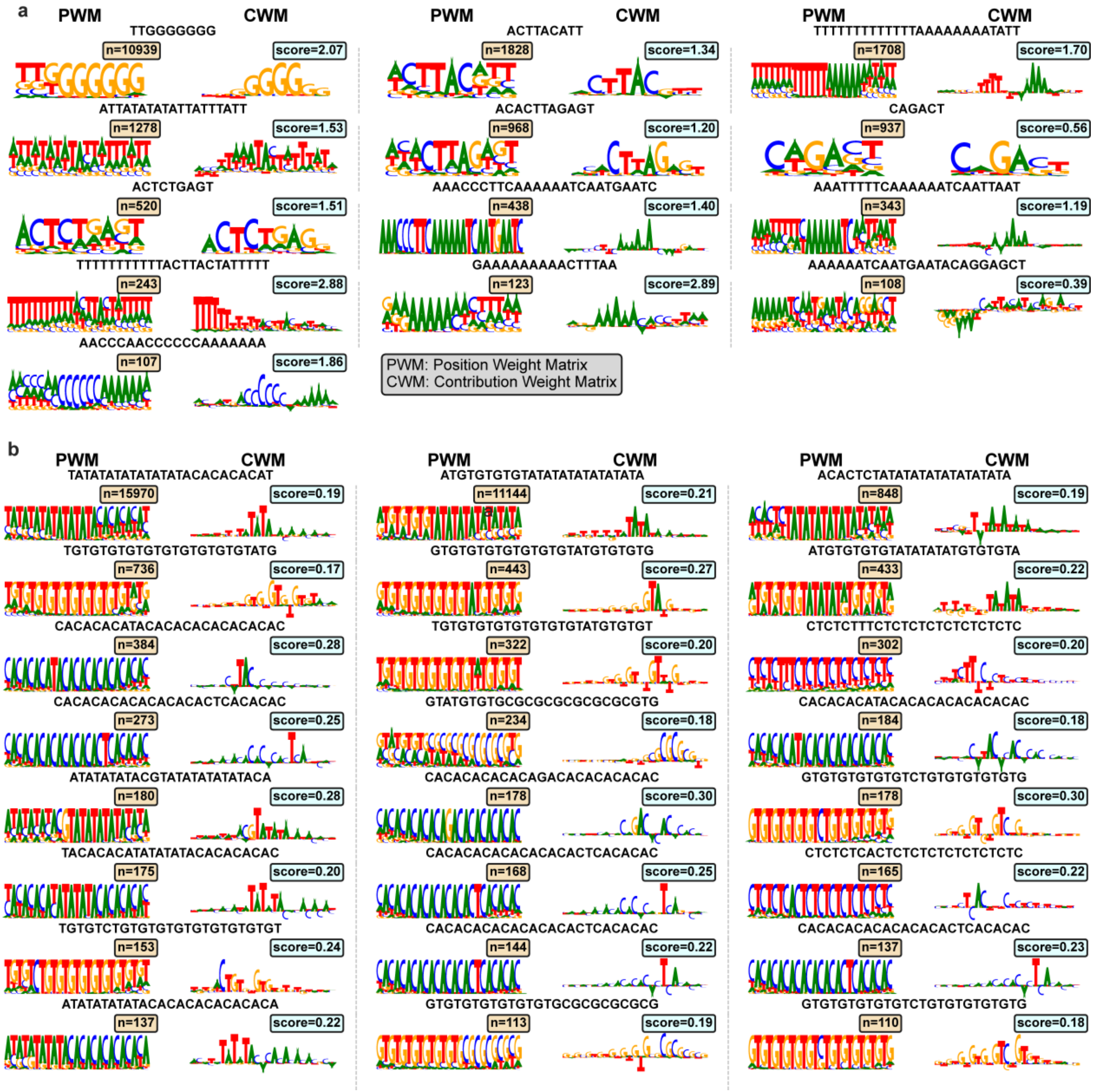


**Supplementary Fig. 19 Position weight matrices (PWMs) and contribution weight matrices (CWMs) of 1-bp and 2-bp insertion motifs.** (**a**) Motifs inferred from 1-bp insertion predictions. (**b**) Motifs inferred from 2-bp insertion predictions.

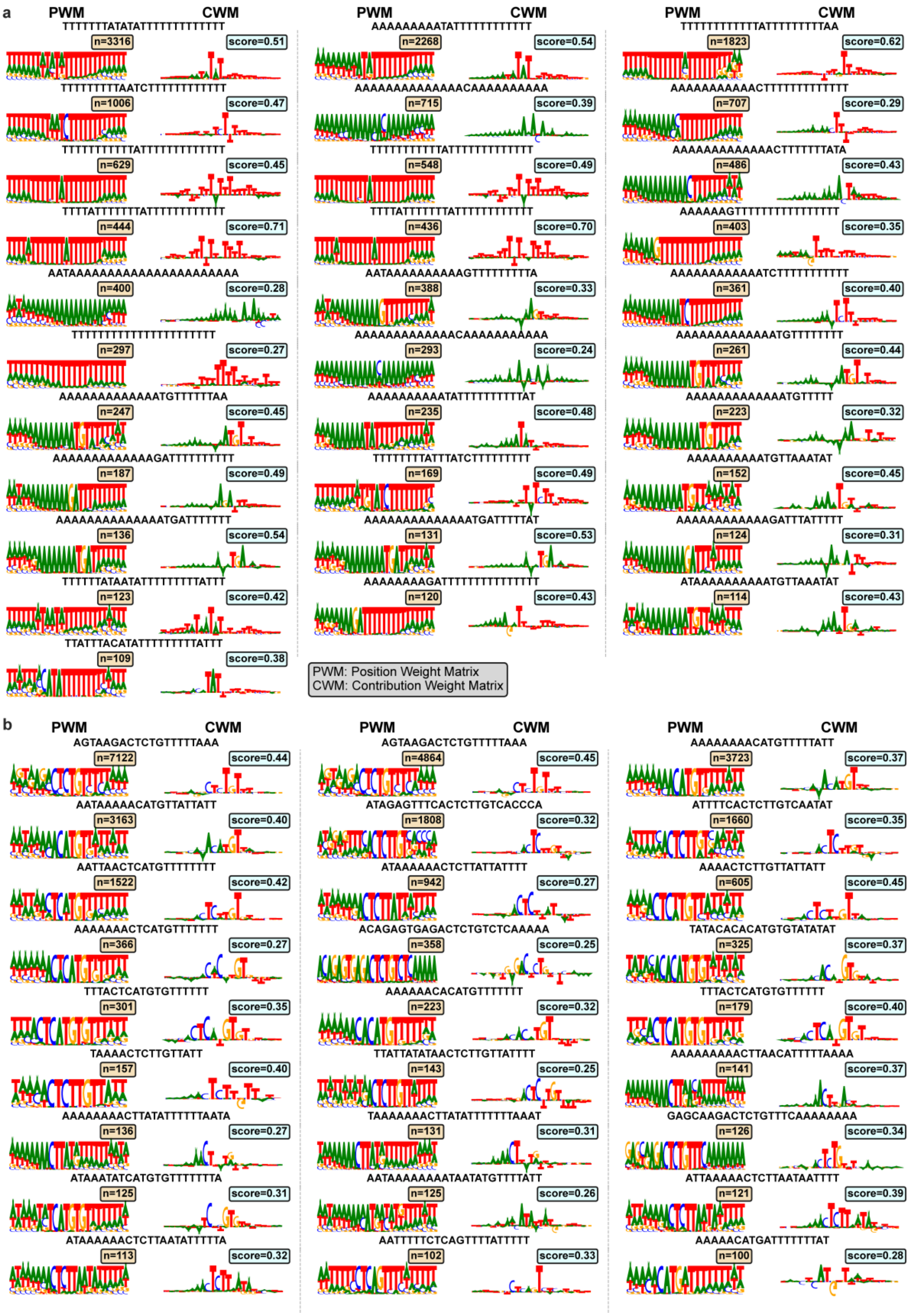


**Supplementary Fig. 20 Position weight matrices (PWMs) and contribution weight matrices (CWMs) of 1-bp and 2-bp deletion start motifs.** (**a**) Motifs inferred from 1-bp deletion start predictions. (**b**) Motifs inferred from 2-bp deletion start predictions.

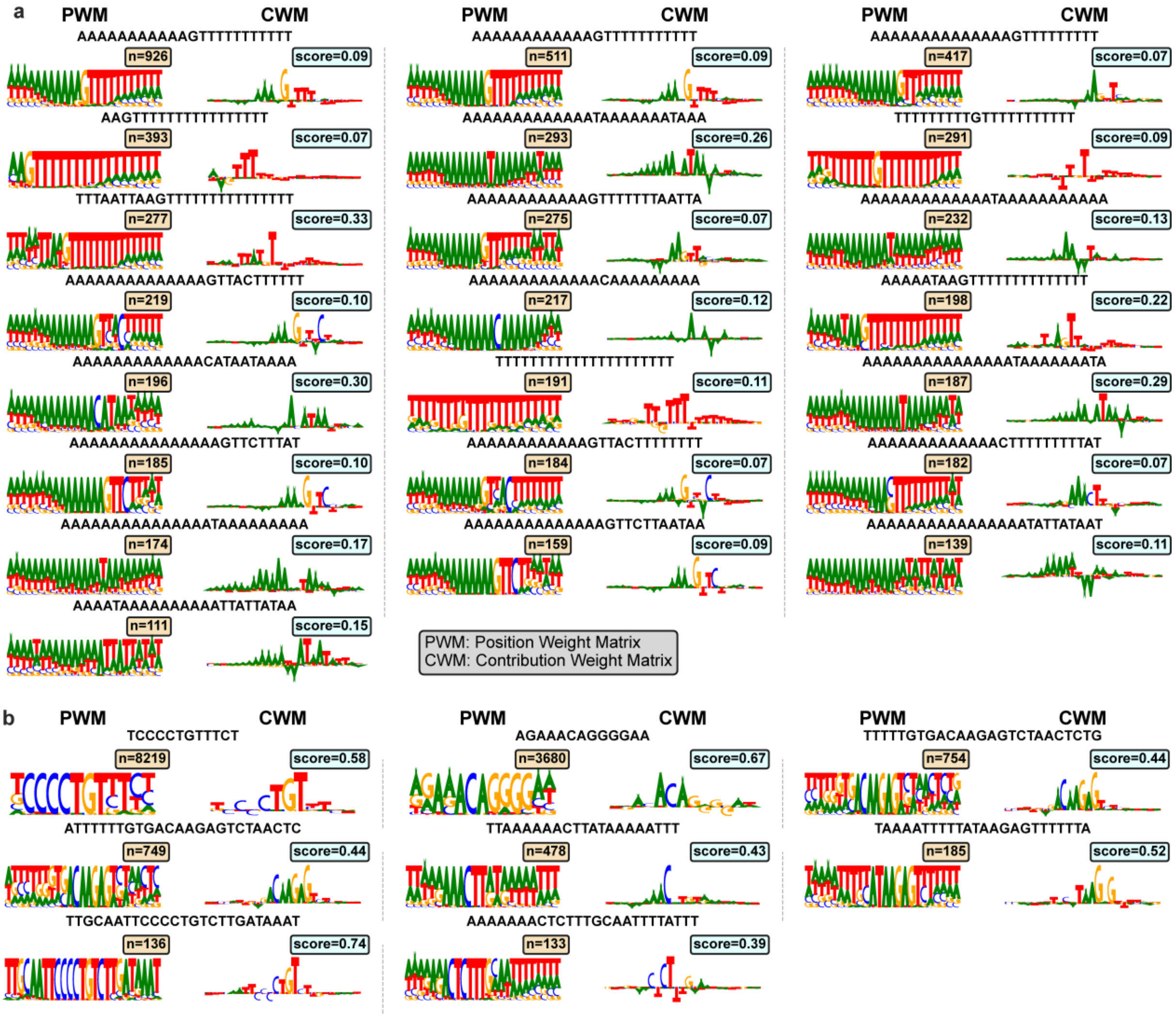


**Supplementary Fig. 21 Position weight matrices (PWMs) and contribution weight matrices (CWMs) of 1-bp and 2-bp deletion end motifs. (a)** Motifs inferred from 1-bp deletion end predictions; **(b)** Motifs inferred from 2-bp deletion end predictions.

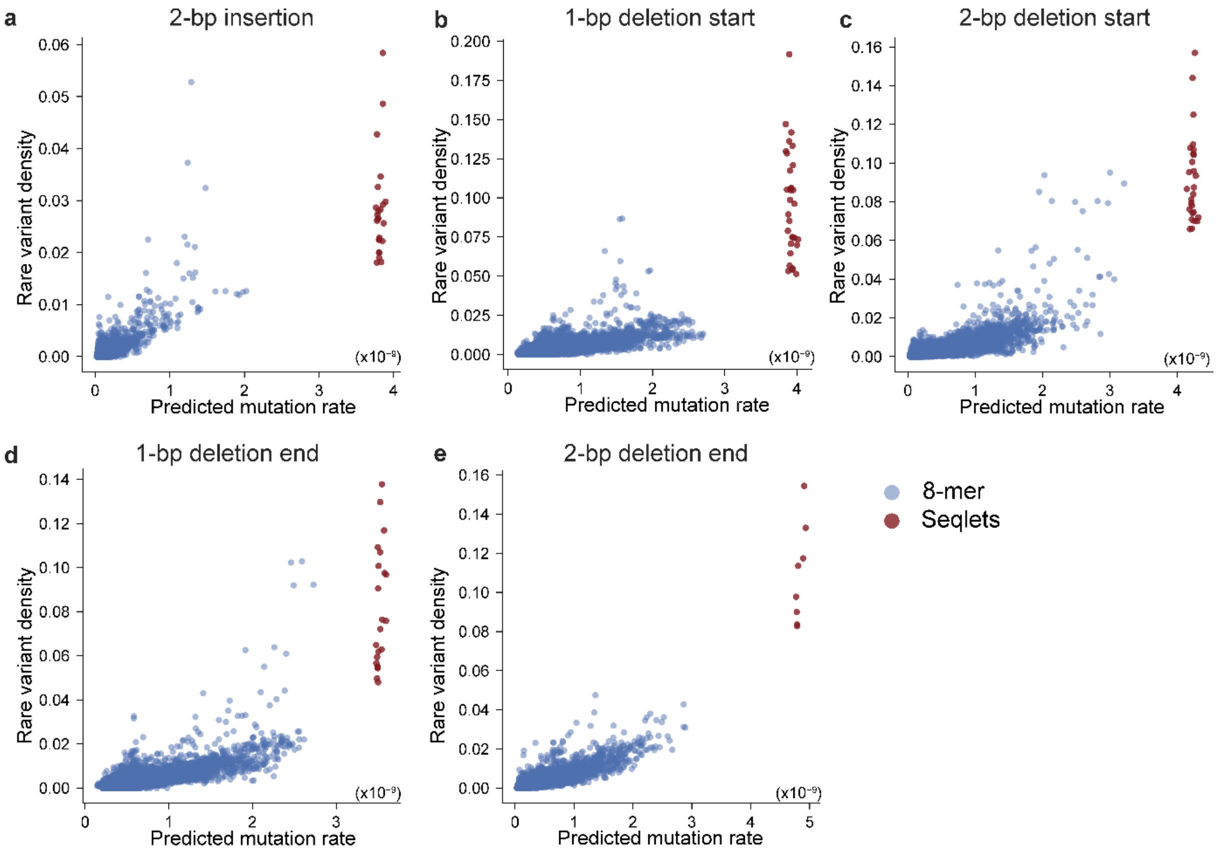


**Supplementary Fig. 22 Comparing rare variant densities and predicted INDEL mutation rates of motifs.** Red dots represent the identified motifs (seqlets) based on central 200bp regions of hypermutated sites; blue dots represent all possible 8-mer motifs.

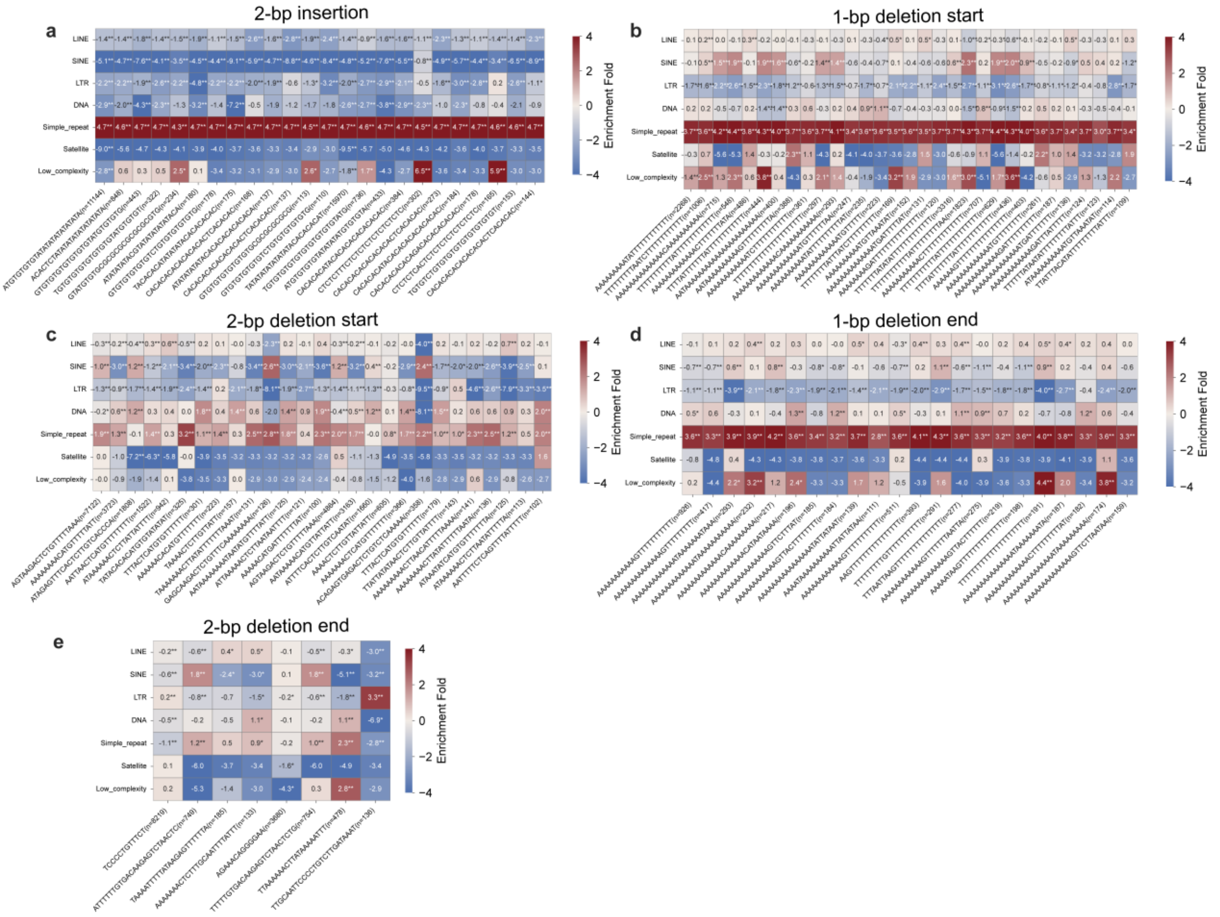


**Supplementary Fig. 23 Enrichment analysis of 1-bp and 2-bp INDEL motifs in the repeat classes.**

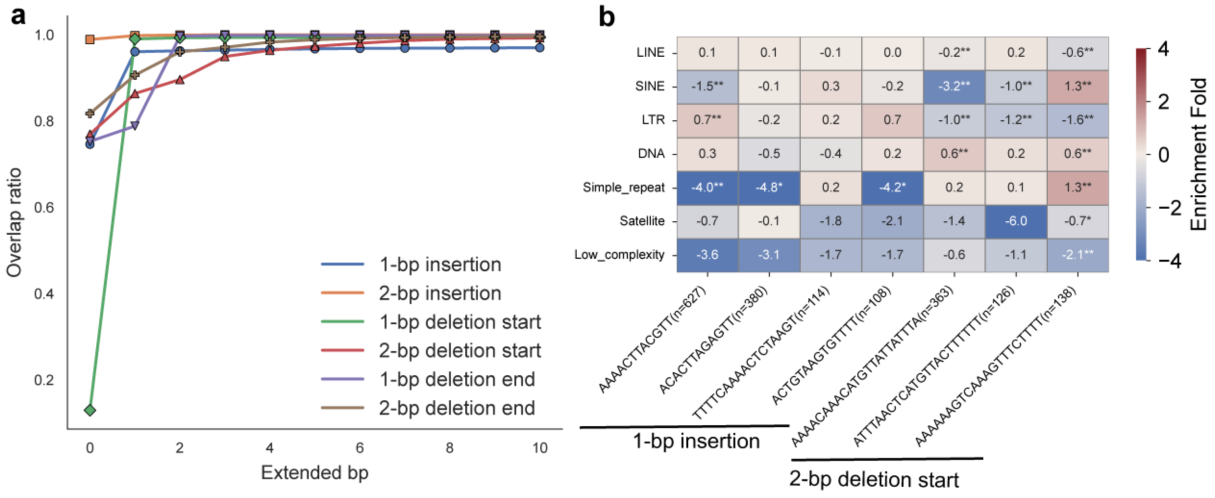


**Supplementary Fig. 24 Motif enrichment analysis in repeats after excluding simple repeats.** (**a**) Fraction of top 100,000 1-bp and 2-bp INDEL sites overlapping simple repeat regions, shown for varying flanking lengths and each INDEL category. X-axis: extension from the central site (bp); Y-axis: percentage of overlapping sites. (**b**) Enrichment of motifs in the repeat classes after filtering hypermutated sites overlapping simple repeats.

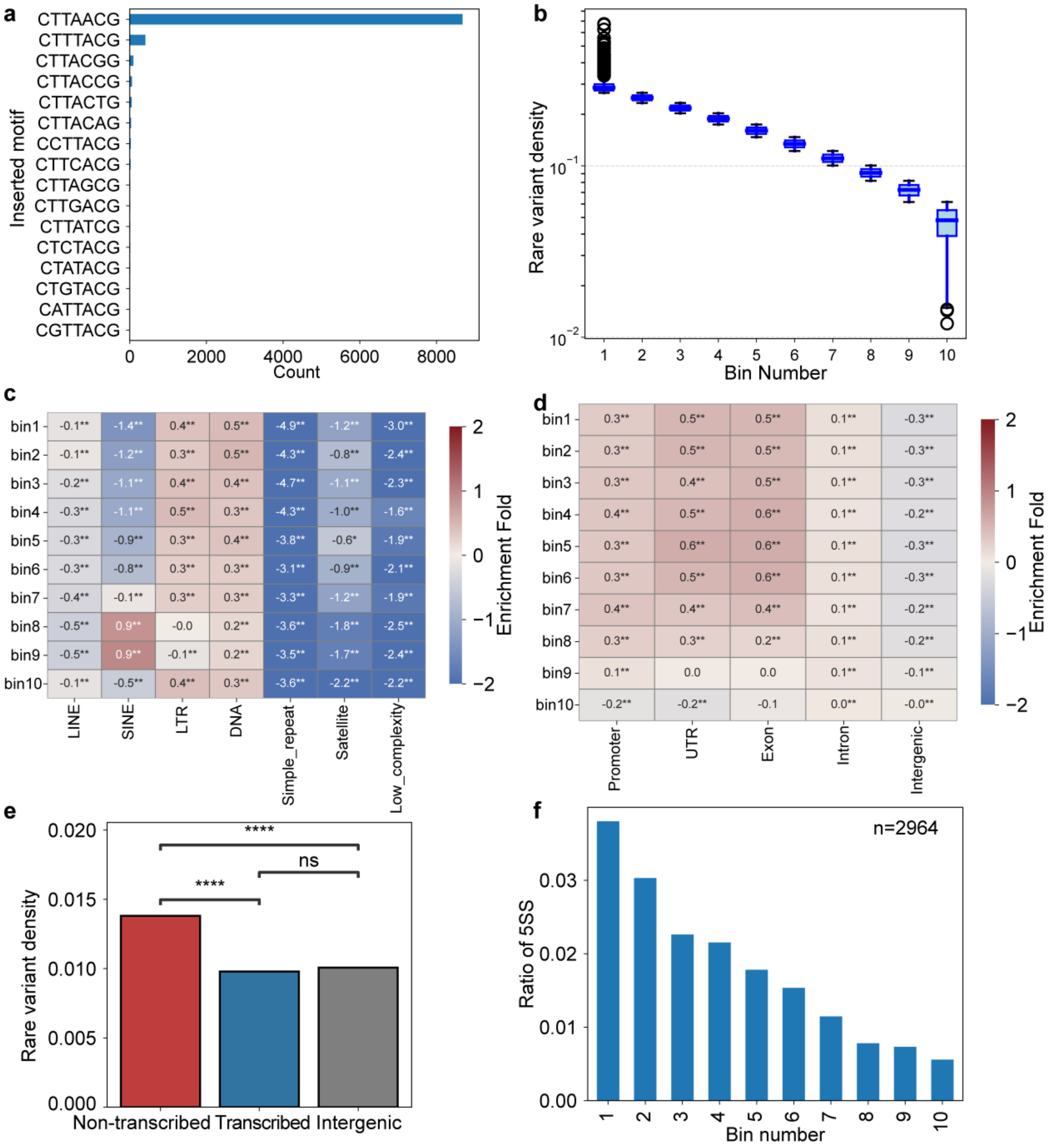


**Supplementary Fig. 25 Association of high 1-bp insertion mutability of the CTTACG motif with transcription and 5′ splice sites.** (**a**) Distribution of CTTACG motifs after 1-bp insertions. (**b**) CTTACG motifs divided into 10 bins based on predicted 1-bp insertion mutation rate quantiles. Y-axis shows mutation rates of a log scale. (**c**) Repeat enrichment analysis for the 10 bins. (**d**) Functional enrichment analysis for the 10 bins. (**e**) Rare variant densities of 1-bp insertions for motifs in transcribed, non-transcribed, and intergenic regions. (**f**) Proportion of motifs containing 5′ splice sites (5SS) in each bin.

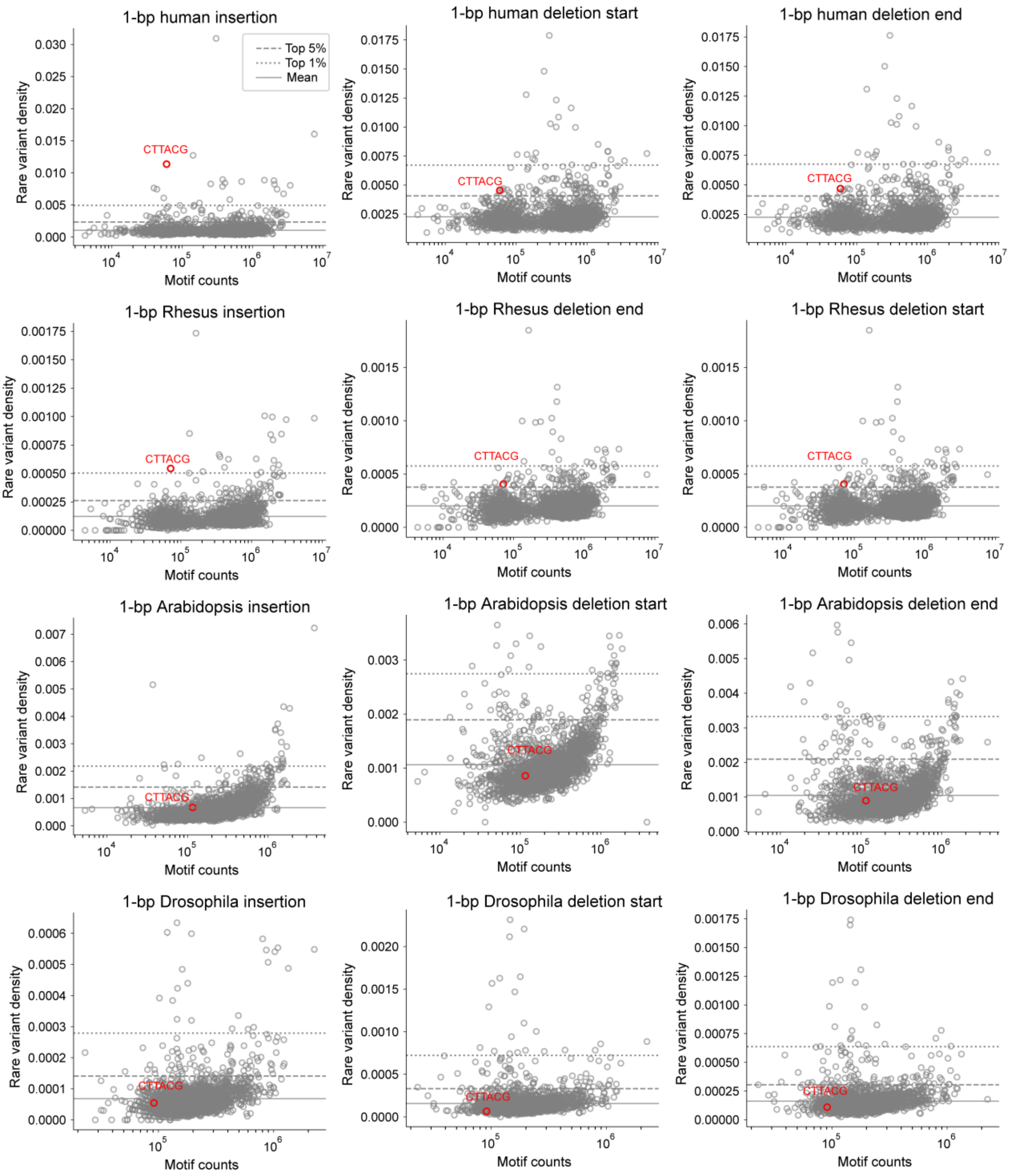


**Supplementary Fig. 26 Rare variant densities of 1-bp insertions at the CTTACG motif across multiple species.** In humans and *M. mulatta*, the rare variant density of 1-bp insertions at the CTTACG motifs is among the top 1% of all possible 6-mer motifs, while that of 1-bp deletions is among the top 5%. In *A. thaliana* and *D. melanogaster*, the rare variant density of 1-bp insertions at the CTTACG motifs is close to or lower than the genome-wide mean. The solid and dotted lines indicate the levels of mutability.

**Supplementary Table 1 Basic statistics for *de novo* INDELs and low frequency INDELs.**

| | **Deletion** | **Insertion** | **Total** |
|---|---|---|---|
| 1in200 | 1,094,980 | 1,112,390 | 2,207,370 |
| 1in2000 | 2,490,212 | 2,193,993 | 4,684,205 |
| 1in20000 | 7,484,568 | 5,236,876 | 12,721,444 |
| 1in100000 | 15,130,208 | 8,124,721 | 23,254,929 |
| *De novo* INDEL | 30,220 | 16,403 | 46,623 |

**Supplementary Table 2 Numbers of training and validation mutations for human INDEL models.**

| **Insertion (1in100000)** | **Training** | **Validation** |
|---|---|---|
| **1bp** | 524,940 | 52,605 |
| **2bp** | 135,705 | 13,394 |
| **3bp** | 70,052 | 6,974 |
| **4bp** | 81,363 | 8,017 |
| **5bp** | 31,238 | 3,112 |
| **6bp** | 28,823 | 2,960 |
| **7-20bp** | 127,879 | 12,938 |
| **Total** | 1,000,000 | 100,000 |
| **Deletion start (1in100000)** | **Training** | **Validation** |
| **1bp** | 900,953 | 90,020 |
| **2bp** | 333,649 | 33,532 |
| **3bp** | 218,881 | 21,770 |
| **4bp** | 208,215 | 20,663 |
| **5bp** | 86,797 | 8,907 |
| **6bp** | 50,237 | 5,031 |
| **7-20bp** | 201,268 | 20,077 |
| **Total** | 2,000,000 | 200,000 |
| **Deletion end (1in100000)** | **Training** | **Validation** |
| **1bp** | 902,552 | 90,093 |
| **2bp** | 335,276 | 33,597 |
| **3bp** | 218,612 | 21,531 |
| **4bp** | 206,825 | 20,803 |
| **5bp** | 86,397 | 8,660 |
| **6bp** | 50,115 | 5,003 |
| **7-20bp** | 200,223 | 20,313 |
| **Total** | 2,000,000 | 200,000 |

**Supplementary Table 3 Numbers of training and validation mutations for *ab initio* and transfer learning models of *M. mulatta*.**

| **Insertion** | **All mutation** | **Training (*ab initio*)** | **Validation (transfer)** | **Training (*ab initio*)** | **Validation (transfer)** |
|---|---|---|---|---|---|
| **1bp** | 408,937 | 203,788 | 20,450 | 81,231 | 8,241 |
| **2bp** | 135,153 | 67,153 | 6,666 | 26,831 | 2,616 |
| **3bp** | 80,724 | 39,969 | 4,146 | 16,196 | 1,638 |
| **4bp** | 76,447 | 38,045 | 3,797 | 15,176 | 1,509 |
| **5bp** | 50,808 | 25,534 | 2,490 | 10,113 | 1,025 |
| **6bp** | 41,806 | 20,721 | 2,075 | 8,435 | 758 |
| **7-20bp** | 210,371 | 104,790 | 10,376 | 42,018 | 4,213 |
| **total** | 1,004,246 | 500,000 | 50,000 | 200,000 | 20,000 |
| **Deletion start** | **All mutation** | **Training (*ab initio*)** | **Validation (transfer)** | **Training (*ab initio*)** | **Validation (transfer)** |
| **1bp** | 583,763 | 186,183 | 18,568 | 74,604 | 7,529 |
| **2bp** | 285,224 | 91,084 | 8,999 | 36,144 | 3,585 |
| **3bp** | 185,125 | 58,884 | 5,922 | 23,721 | 2,421 |
| **4bp** | 188,943 | 60,287 | 6,056 | 24,161 | 2,404 |
| **5bp** | 85,345 | 27,062 | 2,748 | 10,752 | 1,070 |
| **6bp** | 48,259 | 15,471 | 1,532 | 6,094 | 609 |
| **7-20bp** | 191,761 | 61,029 | 6,175 | 24,524 | 2,382 |
| **total** | 1,568,420 | 500,000 | 50,000 | 200,000 | 20,000 |
| **Deletion end** | **All mutation** | **Training (*ab initio*)** | **Validation (transfer)** | **Training (*ab initio*)** | **Validation (transfer)** |
| **1bp** | 581,936 | 186,847 | 18,639 | 74,546 | 7,497 |
| **2bp** | 283,897 | 90,869 | 9,108 | 36,002 | 3,541 |
| **3bp** | 184,289 | 58,767 | 5,961 | 23,607 | 2,315 |
| **4bp** | 188,244 | 59,940 | 6,028 | 24,354 | 2,496 |
| **5bp** | 84,953 | 27,359 | 2,607 | 10,948 | 1,086 |
| **6bp** | 47,939 | 15,271 | 1,571 | 6,060 | 610 |
| **7-20bp** | 190,655 | 60,947 | 6,086 | 24,483 | 2,455 |
| **total** | 1561913 | 500,000 | 50,000 | 200,000 | 20,000 |

**Supplementary Table 4 INDEL data of *A. thaliana*.**

| | Insertion | Deletion start | Deletion end |
|---|---|---|---|
| All | 163,891 | 238,720 | 237,795 |
| Training | 50,000 | 50,000 | 50,000 |
| Validation | 5,000 | 5,000 | 5,000 |

**Supplementary Table 5 INDEL data of *D. melanogaster*.**

| | **Insertion** | **Deletion start** | **Deletion end** |
|---|---|---|---|
| All | 23,427 | 56,357 | 56,357 |
| Training | 10,000 | 20,000 | 20,000 |
| Validation | 1,000 | 2,000 | 2,000 |

**Supplementary Table 6 Relationship between channel numbers and model sizes for MuRaL-indel models.**

| Models | #channels in U-Net layers | #Parameters |
|---|---|---|
| channel_8 | 8、16、24、32、40、48 | 177,912 |
| channel_16 | 16、32、48、64、80、96 | 705,768 |
| channel_24 | 24、48、72、96、120、148 | 1,583,576 |
| channel_32 | 32、64、96、128、160、192 | 2,811,336 |

**Supplementary Table 7 Configuration of key hyperparameters for training human MuRaL-indel models.**

| Hyperparameter | MuRaL-indel models |
|---|---|
| radius | 4000 |
| downsample stride list | 1、4、5、5、5、2 |
| kernel_size | 7 |
| channels | 8 |
| learning_rate | 0.001 |
| weight_decay | 0.01 |
| weight_decay_auto | 0.01 |
| LR_gamma | 0.98 |

**Supplementary Table 8 Configuration of key hyperparameters for training MuRaL-indel models of *M. mulatta*.**

| Hyperparameter | *Ab initio* models | Transfer learning models |
|---|---|---|
| radius | 4000 | 4000 |
| downsample stride list | 1、4、5、5、5、2 | 1、4、5、5、5、2 |
| kernel_size | 7 | 7 |
| channels | 8 | 8 |
| learning_rate | 0.001 | 0.001 |
| weight_decay | 0.01 | 0.01 |
| weight_decay_auto | 0.01 | 0.01 |
| LR_gamma | 0.98 | 0.98 |

**Supplementary Table 9 Configuration of key hyperparameters for training MuRaL-indel models of *A. thaliana*.**

| Hyperparameter | Insertion / deletion start / deletion end model |
|---|---|
| radius | 2000 |
| downsample stride list | 1、4、5、5、5、2 |
| kernel_size | 7 |
| channels | 8 |
| learning_rate | 0.001 |
| weight_decay | 0.01 |
| weight_decay_auto | 0.01 |
| LR_gamma | 0.98 |

**Supplementary Table 10 Configuration of key hyperparameters for training MuRaL-indel models of *D. melanogaster*.**

| Hyperparameter | Insertion / deletion start / deletion end model |
|---|---|
| radius | 2000 |
| downsample stride list | 1、4、5、5、5、2 |
| kernel_size | 7 |
| channels | 8 |
| learning_rate | 0.001 |
| weight_decay | 0.01 |
| weight_decay_auto | 0.01 |
| LR_gamma | 0.98 |
